# The dynamic crosstalk between cytoskeletal filaments regulates the cytoplasmic mechanics across the apicobasal axis

**DOI:** 10.1101/2024.07.23.604784

**Authors:** Dipanjan Ray, Deepak Kumar Sinha

## Abstract

The cytoplasm exhibits viscoelastic properties, displaying both solid and liquid-like behavior, and can actively regulate its mechanical attributes. The cytoskeleton is a major regulator among the numerous factors influencing cytoplasmic mechanics. We explore the interdependence of various cytoskeletal filaments and the impact of their density on cytoplasmic viscoelasticity. The heterogeneous distribution of these filaments give rise to polarised mechanical properties of the cytoplasm along the apicobasal axis. Actin filament (F-actin) disassembly softens the basal cytoplasm while stiffening the mid-cytoplasm due to increased vimentin filament assembly. Disruption of microtubules (MT) or depletion of vimentin softens both the basal and mid-cytoplasm. Cyto D treatment results in localised increase of vimentin assembly in the mid cytoplasm which is dependent on the cytolinker plectin. Nocodazole treatment has a negligible effect on F-actin distribution but significantly alters vimentin’s spatial arrangement. We demonstrate that Cyto D treatment upregulates vimentin expression via ROS-mediated activation of NF-κΒ. This manuscript investigates how different cytoskeletal filaments influence the rheological characteristics of various cytoplasmic regions.

## Introduction

The cytoplasm is a gel-like substance enclosed within the cell membrane that plays a critical role in maintaining cellular structure and facilitating various biochemical processes (Fels et al., 2009; Pollard, 1976; Trevors, 2011). It exhibits viscoelastic properties, possessing both viscous and elastic characteristics that enable it to function effectively (Moeendarbary et al., 2013). This unique nature allows the cytoplasm to behave like a fluid under slow deformations, permitting the smooth movement and distribution of organelles and molecules (Hu et al., 2017). Conversely, it acts like an elastic solid when subjected to rapid stress (Munder et al., 2016). This viscoelasticity is crucial for cellular processes such as motility (Bray, 2000), cell division (Xie et al., 2022), and intracellular transport (Guigas et al., 2007), providing the necessary resilience and flexibility. The viscoelastic properties of the cytosol are influenced by several factors, including the composition and organization of the cytoskeleton (Bausch et al., 1998), the concentration of cytoplasmic proteins (Ellis, 2001; Zimmerman & Minton, 1993), and the presence of cellular organelles (Najafi et al., 2023). The cytoskeleton, consisting of actin, microtubules, and intermediate filaments, provides structural support that enhances the cytoplasm’s elasticity and its capacity to withstand mechanical stress (Pegoraro et al., 2017). The density and cross-linking of these filaments directly affect the cytoplasm’s mechanical properties (Pegoraro et al., 2017). The density and distribution of cytoskeletal filaments are heterogeneous across the cytoplasm. For example, actin filaments typically form a dense meshwork at the cell periphery, offering structural support and aiding in cell shape maintenance (Svitkina, 2020). In contrast, in the cell interior, actin filaments may organize into stress fibers, facilitating cell contraction and motility (Tojkander et al., 2012). Similarly, microtubule filaments exhibit a dynamic distribution pattern throughout the cytoplasm, extending from the microtubule organizing centre (MTOC) towards the cell periphery (Sanchez & Feldman, 2017; Schuh & Ellenberg, 2007). Vimentin, a type III intermediate filament protein, is essential for maintaining cellular integrity, providing mechanical strength, and supporting cellular structures (Liu et al., 2015). Vimentin filaments form a network that resists mechanical stress (Mendez et al., 2014). These filaments create a dynamic network within the cytoplasm, enveloping the nucleus and extending throughout the cell to provide structural support and organization (Zhang et al., 2021). Studies have shown that de novo assembly of vimentin filaments throughout the cytoplasm significantly contributes to the cytoskeletal architecture (Sarria et al., 1990). These cytoskeletal components can interact either directly or via cytoskeletal cross-linkers such as plectin (Wiche, 1998). The interactions between these filaments create a dynamic, meshwork-like pattern within the cytoplasm, which can reorganise in response to various internal and external stimuli (Hohmann & Dehghani, 2019). The distribution of these cytoskeletal filaments varies across different regions of a cell, leading to significant heterogeneity in the cytoplasmic mechanical properties. Several studies have been conducted on the mechanical properties of the cytoplasm, but most provide information about the overall mechanical behaviour without offering much detail about the localised viscoelastic properties of different cytoplasmic regions. The rheology of cytoskeletal filaments has been extensively characterised in various in vitro setups; however, their behaviour inside the cytoplasm is not well understood and may differ significantly. Additionally, how the density and crosstalk between the cytoskeletal filaments affect cytoplasmic mechanics across the apicobasal axis remains elusive. This study reports that the density of actin, microtubule, and vimentin filaments is anisotropic along the apicobasal axis of NIH 3T3 fibroblasts. Using particle tracking microrheology (PTM), we demonstrate that the mechanical properties of the cytoplasm along the apicobasal axis are directly influenced by the density of these cytoskeletal filaments. This manuscript establishes the role of individual cytoskeletal filaments in determining the local mechanical properties of the basal and mid-cytoplasm. Furthermore, our findings indicate that crosstalk between cytoskeletal components is crucial for maintaining the mechanical integrity of the cytoplasm. Additionally, we report that the actin-disrupting agent Cyto D enhances the expression of vimentin via ROS-dependent activation of NF-κB.

## Results

### Mechanical properties of the cytoplasm are polarised along the apicobasal axis

The cytoplasm contains various cytoskeletal components, such as actin microfilaments, microtubules (MT) and intermediate filaments (IF) (Hohmann & Dehghani, 2019). However, their distribution and abundance are not uniform and vary along the apicobasal axis. **Figure 1A** illustrates the distribution of these three major cytoskeletal filaments: actin, microtubules, and vimentin (a type III intermediate filament). This heterogeneous distribution results in locally variable mechanical properties of the cytosol (Fletcher & Mullins, 2010). To evaluate these local mechanical properties, we analysed the trajectories (r(t)) of fluorescently labelled microspheres, each with a diameter of 200 nm, scattered throughout the cytoplasm (**Figure 1B**). The trajectories were derived by monitoring the random, spontaneous movement of the microsphere centroids across 1000 frames over a 20-second duration, corresponding to one frame every 20 milliseconds. These microspheres serve as probes for the local viscoelastic properties of the cytoplasm. The mean squared displacement (MSD) values obtained from the trajectories of these microspheres provide insight into the local viscoelastic properties of the cytoplasm. The time-scale-dependent compliance values derived from MSDs describe the local deformability of the cytoplasm in response to thermal forces ∼ kBT/a. Here, kB, T, and a represent Boltzmann’s constant, the absolute temperature of the medium, and the radius of the microsphere, respectively. These forces arise from the thermal fluctuations of the probe microspheres (Wirtz, 2009). **Figure 1C** compares MSDs and compliance of the most and least compliant cytoplasmic regions at ten different locations within a given cell. Consistent with previous findings (Tseng et al., 2002), our results suggest that local deformability varies significantly throughout the cytoplasm of a cell. It is presumed that the average density of cytoskeletal filaments varies along the apicobasal axis of the cytoplasm (**Figure 1A**). Therefore, we hypothesised that these variations in cytoskeletal filaments contributed to the polarisation of the locally heterogeneous mechanical properties of the cytoplasm along the apicobasal axis. To resolve the mechanical properties along the apicobasal axis, regions within 0 < z < 2 microns and z > 2 microns are designated as the basal and mid / apical regions of the cytoplasm, respectively (**Figure 1A**). We designated the first sharply focused DIC image of the edge of the cell as the basal plane during particle tracking microrheology (PTM), while when calculating the fractal dimension (FD) values, the basal plane was determined based on the presence of well-focused actin stress fibres (Maninová & Vomastek, 2016). To minimise the ambiguity that arises from height variations between different cells, we selected cells that had comparable heights (∼4±1μm). A histogram of the cell heights used for analysis is presented in **Figure S1-A**. By analysing the trajectories of at least 60 microspheres embedded in the basal or mid-cytoplasmic regions of at least 10 different cells (**Figure S1-B** and **S1-C**), we determined the mechanical properties of the cytoplasm along the apicobasal axis. **Figure 1D** compares the average MSDs and compliance values of microspheres distributed in the mid and basal cytoplasmic regions of at least 10 different cells. The mean MSDs indicated that the mid cytoplasm demonstrated higher compliance compared to the basal cytoplasm. The frequency-dependent viscoelastic moduli are algebraically derived from these MSD values (Mason, 2000). The dynamic viscosity, η, is calculated by dividing the viscous modulus, G’’, by the corresponding angular frequency, ω. Given the frequency dependence of mechanical properties, we analysed the elastic modulus (G’) and dynamic viscosity (η) at 1 Hz and 10 Hz frequencies to ensure uniformity in comparisons across varying conditions. The elastic modulus (G’) of the mid cytoplasmic region was found to be significantly lower than that of the basal plane at 1Hz and 10Hz, indicating a softer cytoplasm at least 4-5 times (**Figure 1F-i, ii**). The dynamic viscosity (η) of the mid cytoplasmic region remained at least 3 times lower than the basal cytoplasmic region at both frequencies (**Figure 1F-iii, iv**). In conclusion, the mechanical properties of the cytoplasm exhibited polarity along the apicobasal axis, leading to a decrease in both stiffness and viscosity with increasing cell height. When an elastic material undergoes deformation, it stores energy that can be recovered upon stress release. However, in viscoelastic materials like cytoplasm, some of this energy is also dissipated due to microstructural rearrangement, which manifests as the viscous flow of the material. This dissipation of energy is indicated by the dimensionless quantity known as the loss tangent, tan δ, which is the ratio between viscous (G’’) and elastic modulus (G’). A higher value of tan δ indicates that a greater proportion of energy associated with deformation is dissipated as heat. The tan δ also delineates viscoelastic solids from liquids. At the crossover frequency (ω_c_), where the elastic modulus (G’(ω)) intersects with the viscous modulus (G″(ω)), the mechanical properties of the material undergo a transition from a viscoelastic solid (tan δ < 1) to a viscoelastic liquid (tan δ > 1). The reciprocal of ω_c_, designated as the relaxation time (Schwarzl, 1971), represents the timescale over which the stored elastic stresses dissipate due to spontaneous microstructural rearrangements. Given that the loss tangent (tan δ) of the basal and mid cytoplasm remained above 1 across the frequency spectrum of 1-10Hz (**Figure 1E**), it can be inferred that the cytoplasm exhibits mechanical characteristics of a viscoelastic liquid. Furthermore, the decay of tan δ with frequency and an apparent plateau near 10 Hz indicate that the relaxation time for microstructural rearrangements of the cytoplasm is slower than 0.1 sec for both the basal and mid regions of the cytoplasm. Next, we examined the relative density of actin (**Figure 1G-i**), microtubule (**Figure 1G-ii**), and vimentin (**Figure 1G-iii**) filaments across the basal (0 < z < 2μm), mid (2μm < z < 4μm) and apical (z > 4μm) cytoplasmic regions. To quantify the density of cytoskeletal filaments, we computed their fractal dimension (FD) values. The representative binary images of actin, vimentin, and microtubule filaments across the apicobasal axis used for the FD analysis are presented in **Figure S1-D**. **Figure 1H** compares the FD values of different cytoskeletal filaments across the apicobasal axis. The relative abundance of these filaments varies significantly in the basal cytoplasm; however, in the mid and apical regions, the differences in their densities are negligible. Actin and vimentin filaments were predominantly concentrated in the basal cytoplasm, with their densities gradually decreasing towards the mid and apical regions. In contrast, microtubule (MT) filaments showed relatively higher concentration around the mid cytoplasmic region compared to the basal and apical regions. Based on **Figure 1H**, we deduced that disrupting actin filaments would lead to distinct changes in the mechanical properties of the basal and mid cytoplasm. Specifically, we anticipated that both regions would become softer due to actin disassembly.

**Figure 1:**
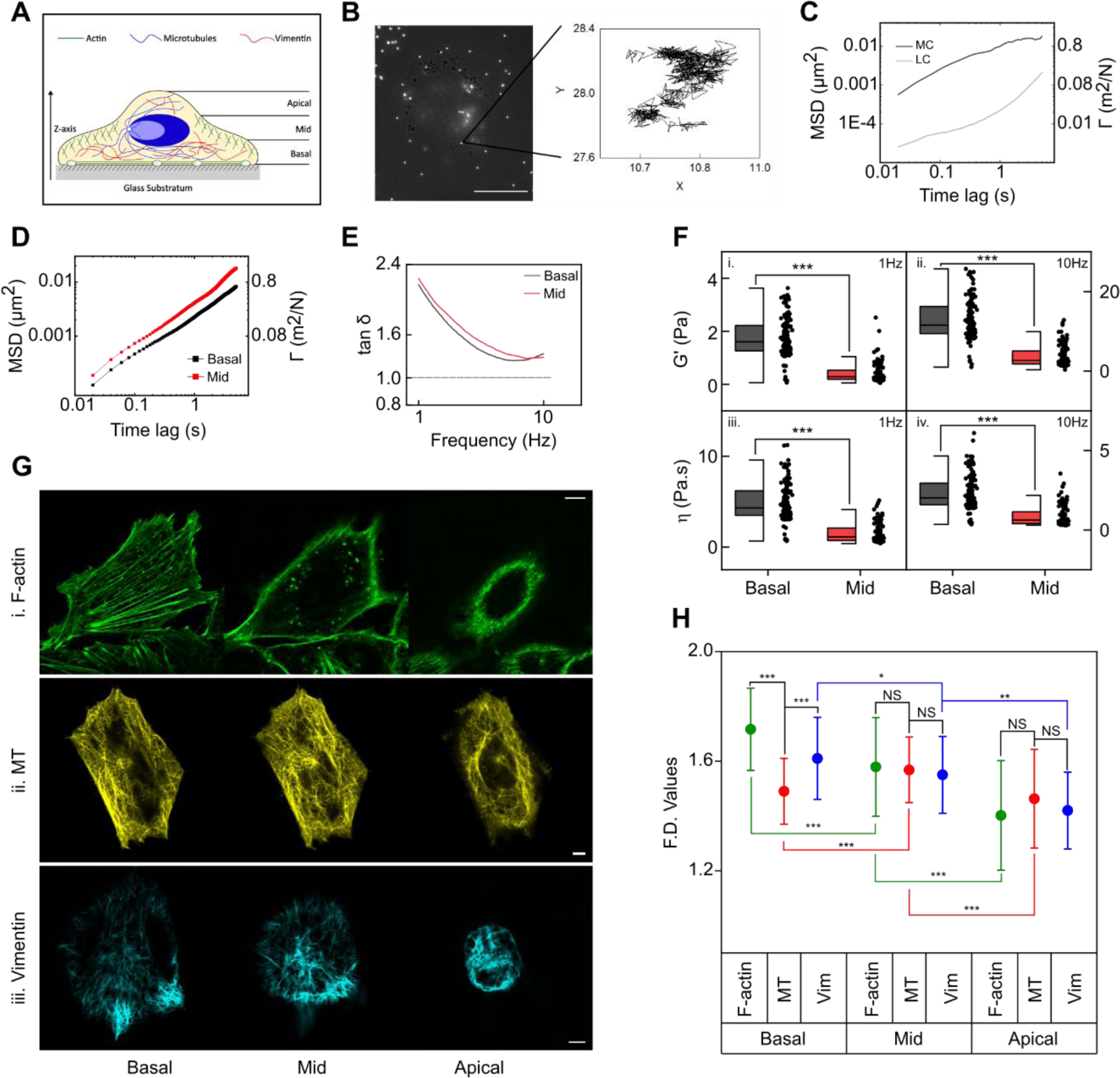
Mechanical properties of the cytoplasm are polarised along the apicobasal axis. (**A**) Cartoon of a cell illustrating the distribution of actin, microtubule, and vimentin filaments, along with segmentation of the cytoplasm into basal, mid, and apical regions along the z-axis. (**B**) Fluorescently labelled microspheres with a diameter of 200 nm dispersed throughout the cytoplasm of an NIH 3T3 fibroblast (Scale bar, 5 μm). (Inset) Trajectory of one such microsphere tracked over a 20-second period. (**C**) MSD (left axis) and compliance, Γ (right axis) of the most compliant (MC) and least compliant (LC) regions within the cytoplasm of a specific cell. (**D**) Mean MSD (left axis) and compliance, Γ (right axis) of the basal and mid cytoplasmic regions. (N_beads_ > 80, N_cells_ > 15) (**E**) Loss tangent profiles of the basal and mid cytoplasmic regions. (**F**) Comparison of elastic modulus (G’) values between the basal and mid cytoplasmic regions at 1 Hz (i) and 10 Hz (ii) frequencies. Comparison of dynamic viscosity (η) values between the basal and mid cytoplasm at 1 Hz (iii) and 10 Hz (iv) frequencies. (N_beads_ > 80, N_cells_ > 15; ***p<0.001, Mann-Whitney U test) (**G**) Representative confocal images of F-actin (i), microtubule, MT (ii) and vimentin filaments (iii) across the basal, mid and apical cytoplasm. (Scale bar, 5 μm) (**H**) Fractal dimension (FD) values of actin, microtubule (MT) and vimentin (Vim) filaments across the apicobasal axis. (Mean ± SD, N_cells_ > 100; *p<0.05, **p<0.01, ***p<0.001, NS, non-significant, two sample T-test)

### Disruption of actin filaments has contrasting effects on the mechanical properties of the basal and mid cytoplasm

Next, using cytochalasin D (Cyto D), we investigated the impact of actin filament disruption on the mechanical properties of the basal and mid cytoplasmic regions. Cyto D inhibits the polymerization of actin by binding to its barbed ends (Brown & Spudich, 1981; Casella et al., 1981; Cooper, 1987; Masiello et al., 1982). **Figure 2A** compares the mechanical compliance, Γ(t) and the MSD, between control and Cyto D treated cells. Disruption of actin filaments by Cyto D reversed the mechanical polarity of the cytoplasm along the apicobasal axis by making the mid cytoplasm less compliant than the basal cytoplasm (**Figure 2A-i**). Comparison with control cells reveals that Cyto D treatment increased the compliance of the basal cytoplasm significantly (**Figure 2A-ii**), while the mid cytoplasm became slightly stiffer (**Figure 2A-iii**). Considering that the basal cytoplasm contained a higher concentration of actin filaments compared to the mid cytoplasm, we anticipated a significantly larger increase in the compliance of the basal plane than the mid cytoplasm due to actin disassembly. However, contrary to the anticipation we observed that actin disassembly caused a relatively smaller decrease of compliance in the mid cytoplasmic region (**Figure 2A-iii**). As anticipated, the disassembly of actin led to greater fluidization of the basal cytoplasm compared to the mid cytoplasm (**Figure S2-A**). Next, we examined the effect of actin disassembly on the elastic modulus (G’) and dynamic viscosity (η) of the basal and mid cytoplasm. A study by Rotsch & Radmacher (2000) reported that disassembly of actin filaments led to a significant decrease in the cell’s elastic modulus, highlighting the critical role of the actin network in maintaining cellular mechanical stability. Similar to it, we found that the median elastic modulus (G’) of the basal cytoplasm decreased significantly, by 8-10 fold at both 1 Hz and 10 Hz frequencies, due to the disassembly of actin filaments by cytochalasin D (**Figure 2B-i,ii**). Similarly, the dynamic viscosity (η) of the basal cytoplasm decreased 7-9 fold as a result of Cyto D treatment (**Figure 2B-iii,iv**). However, the effect of actin disruption on the mechanical properties (G’ and η) of the mid cytoplasm is contrary to expectations (**Figure 2C**). Cyto D treatment caused a significant increase in both the elastic modulus (G’) (**Figure 2C-i,ii**) and the dynamic viscosity (η) (**Figure 2C-iii,iv**) in the mid cytoplasm. To understand the reason behind the stiffening of the mid cytoplasm, we compared the density of actin filaments between control and Cyto D treated cells. Cyto D treatment (2 μM, 1 hour) resulted in the fragmentation of longer actin filaments into smaller segments (**Figure 2E-i**). Cyto D mediated disassembly of actin caused significant loss of actin filaments in the basal, mid and apical cytoplasm (**Figure 2D-i**). Therefore, we hypothesised that the observed elevation in stiffness, as depicted in **Figure 2C**, despite the loss of actin filaments, was due to the increased assembly of other cytoskeletal filaments within the mid cytoplasmic region. The frequency dependence of the mid cytoplasm’s loss tangent further indicates that there are additional changes in the microstructure of the cytoplasm other than the loss of actin filaments (**Figure S2-A** and **S2-C**). Next, we explored the impact of actin disassembly on the density of microtubule and vimentin filaments. Unlike actin filaments, Cyto D treatment did not have a significant effect on the density of MT filaments (**Figure 2E-ii**). Analysis of FD values indicate that there was no significant change in the density of MT filaments across the apicobasal axis due to Cyto D treatment (**Figure 2D-ii**). Hence, MT filaments cannot be attributed to the stiffening of the mid cytoplasm following Cyto D-induced disruption of actin filaments. **Figure 1H** shows that the density of vimentin filaments was highest at the basal cytoplasm and decreased gradually at the mid and apical cytoplasmic regions. Treatment with Cyto D altered the distribution of vimentin filaments across the apicobasal axis **(Figure 2D-iii)**. While the density of vimentin filaments showed no significant changes in the basal and apical cytoplasm, the mid cytoplasm exhibited a significant increase in the density of vimentin filaments in response to Cyto D treatment. Confocal images clearly show the increased assembly of vimentin filaments in the mid cytoplasm (**Figure 2E-iii**) of Cyto D treated cells. These observations aligned with the observed increase in the elastic modulus (G’) values of the mid cytoplasm of Cyto D treated cells (**Figure 2C-i, ii**). Hence, we hypothesised that the elevated assembly of vimentin filaments was responsible for the increased stiffness of the mid cytoplasm in Cyto D treated cells.

**Figure 2:**
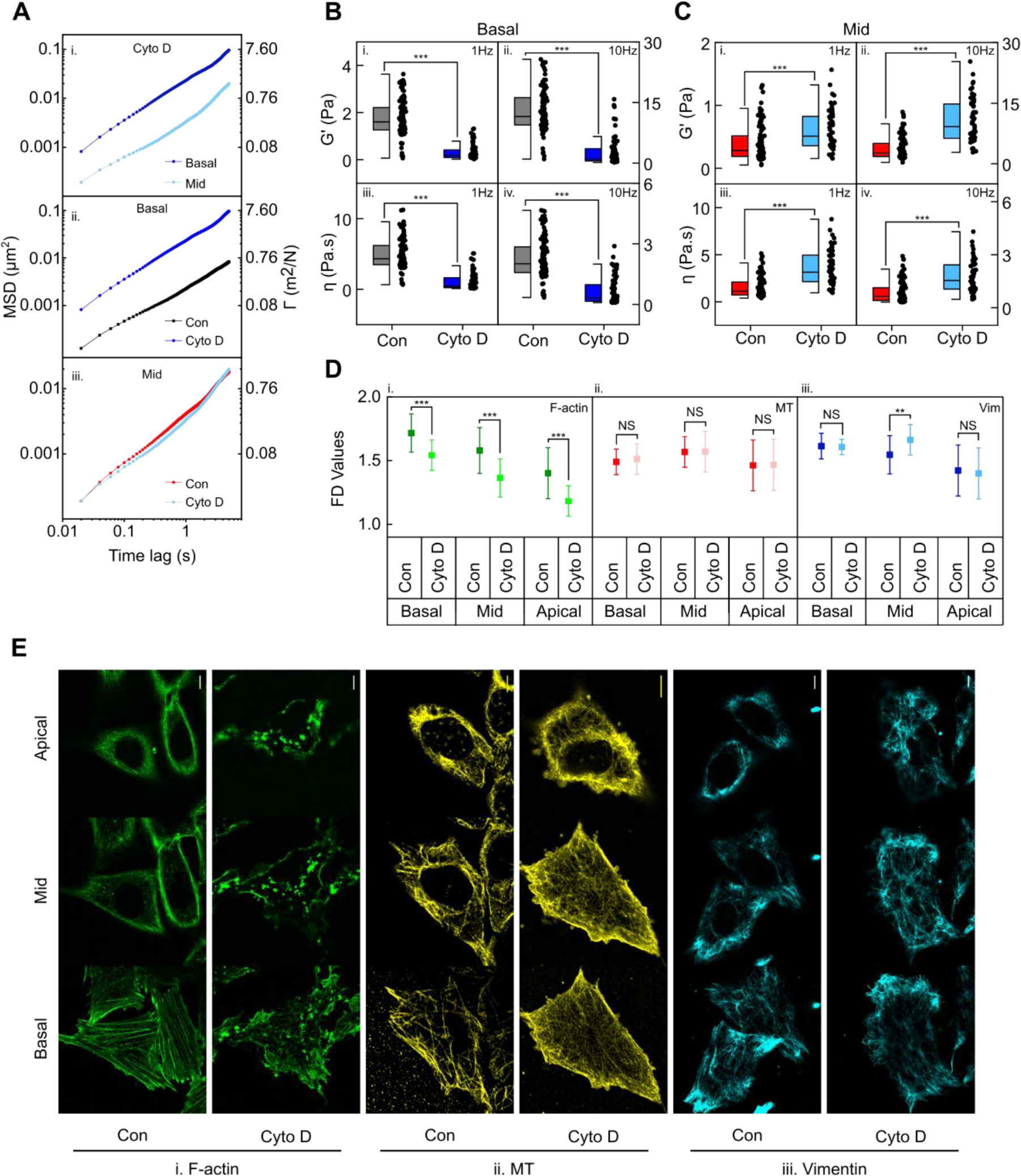
Disruption of actin filaments has contrasting effects on the mechanical properties of the basal and mid cytoplasm. (**A**) Mean MSD (left axis) and compliance, Γ (right axis) of the basal and mid cytoplasmic regions of Cyto D treated cells (i), comparison of Mean MSDs (left axis) and compliance, Γ (right axis) between the control and Cyto D treated cells across the basal (ii) and mid (iii) cytoplasmic regions. (N_beads_ > 80, N_cells_ > 15) (**B**) Comparison of elastic moduli (G’) between the control and Cyto D treated cells across the basal cytoplasmic region at 1 Hz (i) and 10 Hz (ii) frequencies. Comparison of dynamic viscosity (η) values between the control and Cyto D treated cells across the basal cytoplasmic region at 1 Hz (iii) and 10 Hz (iv) frequencies. (N_beads_ > 80, N_cells_ > 15; ***p<0.001, Mann-Whitney U test) (**C**) Comparison of elastic moduli (G’) between the control and Cyto D treated cells across the mid cytoplasmic region at 1 Hz (i) and 10 Hz (ii) frequencies. Comparison of dynamic viscosity (η) values between the control and Cyto D treated cells across the mid cytoplasmic region at 1 Hz (iii) and 10 Hz (iv) frequencies. (N_beads_ > 80, N_cells_ > 15; ***p<0.001, Mann-Whitney U test) (**D**) Fractal dimension (FD) values of actin, microtubule (MT) and vimentin filaments in control and Cyto D treated cells across the apicobasal axis. (Mean ± SD, N_cells_ > 100; **p<0.01, ***p<0.001, NS, non-significant, two sample T-test) (**E**) Representative confocal images of F-actin, microtubules (MT) and vimentin filaments in the control and Cyto D treated cells across the apicobasal axis. (Scale bar, 5 μm)

### Nocodazole-induced depolymerization of microtubules results in significant softening of the cytoplasm across the apicobasal axis

Microtubules, like actin filaments, form intricate networks within the cytoplasm; however, their spatial arrangement significantly differs from that of actin filaments along the apicobasal axis (**Figure 1G-ii**, **Figure 1H**). Consequently, microtubule disassembly is expected to affect the mechanical properties of the cytoplasm differently from the disruption of actin filaments. We investigated the impact of microtubule disassembly on the mechanical properties of the basal and mid cytoplasmic regions. We used nocodazole to disrupt microtubules, which acts by interfering with their polymerisation (Xu et al., 2002). The effect of microtubule disassembly on the compliance of these cytoplasmic regions differed notably from that of actin filament perturbation. Unlike Cyto D treatment, the nocodazole treated cells retained the viscoelastic polarity along the apicobasal axis. The mid cytoplasm of nocodazole treated cells exhibited higher compliance compared to the basal cytoplasm (**Figure 3A-i**). A comparison between **Figure 3A-i** and **Figure 1D** demonstrated that the difference in compliance along the apicobasal axis increased by approximately 7-fold at 1 Hz and 9-fold at 10 Hz due to microtubule disassembly (**Figure S3-A**). The disassembly of microtubules led to a significant increase in compliance in both the basal (**Figure 3A-ii**) and mid cytoplasmic (**Figure 3A-iii**) regions compared to their counterparts in control cells. According to our observations, microtubule disruption had a pronounced effect on the compliance of the mid-cytoplasm compared to the basal cytoplasm. A 4 to 7 fold greater increase in compliance in the mid cytoplasm (**Figure S3-B**) further validated the higher density of microtubules within the mid cytoplasmic region compared to the basal region, as shown in **Figure 1H**. Interestingly the disassembly of microtubule makes the viscosity and G’ of the cytoplasm more polarised along the apicobasal axis (**Figure S3-E**). In the mid cytoplasm, the disassembly of microtubules caused an intriguing change in its mechanical properties. The mid-cytoplasm became more fluid-like at 10 Hz, while it exhibited more gel-like characteristics at lower frequencies (**Figure S3-C**). The frequency dependence of the mid-cytoplasm’s loss tangent further indicated that there were additional changes in the cytoplasm’s microstructure other than the loss of microtubule (**Figure S3-D**). Next, we probed the effect of Nocodazole treatment on the other cytoskeletal filaments. Nocodazole treatment had an insignificant effect on the FD of the actin filaments (**Figure 3D-i**, **Figure 3E-i**). As expected nocodazole treatment (10uM, 1hr) severely altered the MT organization throughout the apicobasal axis by reducing the density of the MT filaments (**Figure 3D-ii**, **Figure 3E-ii**). The global loss of MT filaments upon nocodazole treatment justified the reduction in cytoplasmic stiffness along the entire apicobasal axis. The MT disassembly had a pronounced impact on the vimentin filament network (**Figure 3D-iii, Figure 3E-iii**). We observed a significant reduction of vimentin filaments across the entire cytoplasm (**Figure 3D-iii**). Most of the vimentin filaments were aggregated as thick strands localised in the perinuclear region (**Figure 3E-iii**). Nocodazole treatment altered the relative spatial distribution of vimentin filaments within the cell cytoplasm. In control cells, we observed a decreasing vimentin filament density along the apicobasal axis (**Figure 1H**). Conversely, in nocodazole-treated cells, the vimentin filament density increased along the apicobasal axis (**Figure 3D-iii**). It is difficult to separate the contribution of individual cytoskeletal filaments in governing the mechanical behaviour of the cytoplasm. The basal cytoplasm of the nocodazole treated cells provides ideal system to explore the mechanical characteristics of actin filaments in cells. The global loss of vimentin filaments upon MT disassembly further contributed to the softening of the basal and mid cytoplasmic regions.

**Figure 3:**
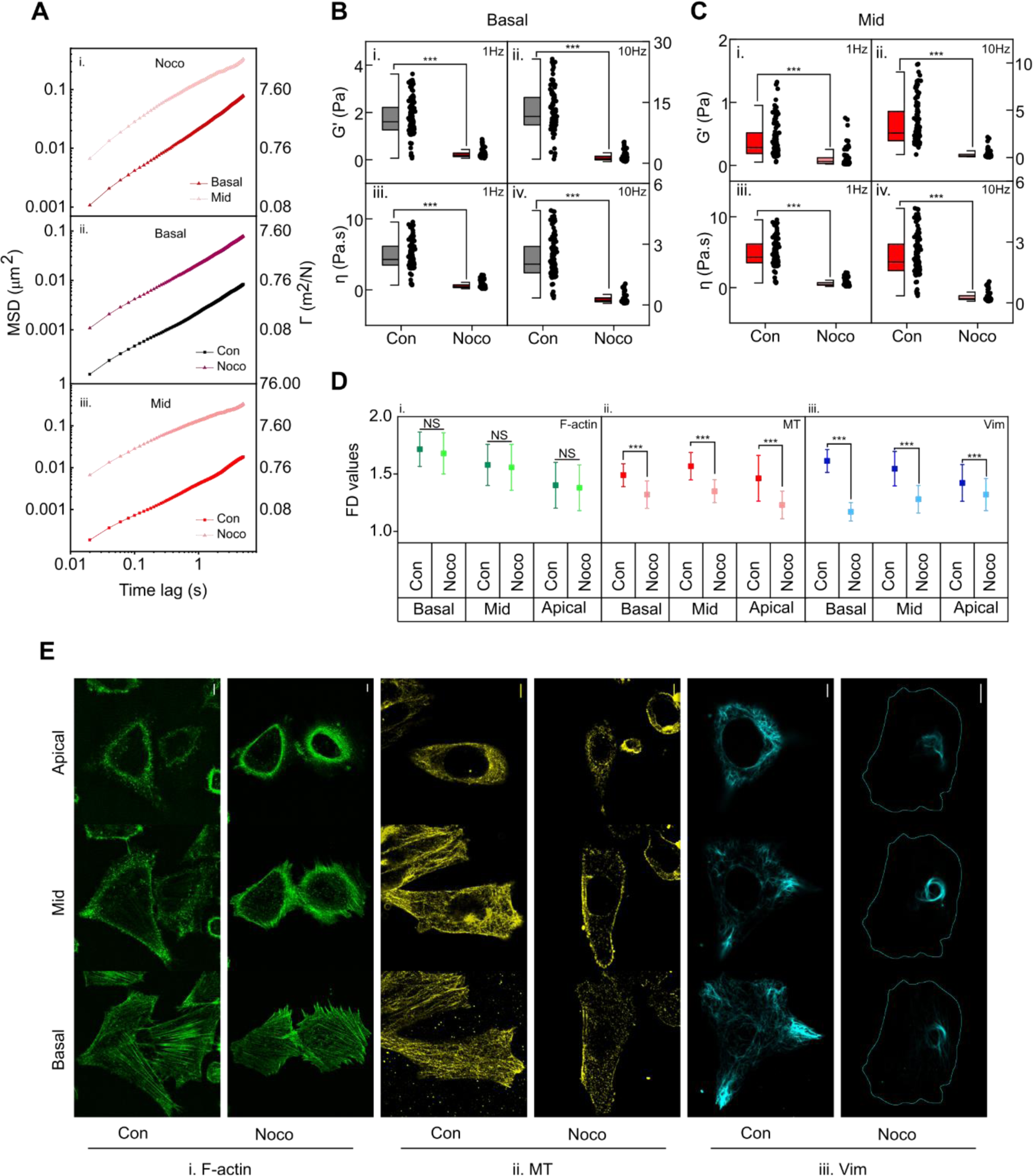
Nocodazole-induced depolymerization of microtubules results in significant softening of the cytoplasm across the apicobasal axis. (**A**) Mean MSD (left axis) and compliance, Γ (right axis) of the basal and mid cytoplasmic regions of nocodazole (noco) treated cells (i), comparison of Mean MSDs (left axis) and compliance, Γ (right axis) between the control and nocodazole treated cells across the basal (ii) and mid (iii) cytoplasmic regions. (N_beads_ > 60, N_cells_ > 10) (**B**) Comparison of elastic moduli (G’) between the control and nocodazole treated cells across the basal cytoplasmic region at 1 Hz (i) and 10 Hz (ii) frequencies. Comparison of dynamic viscosity (η) values between the control and nocodazole treated cells across the basal cytoplasmic region at 1 Hz (iii) and 10 Hz (iv) frequencies. (N_beads_ > 60, N_cells_ > 10; ***p<0.001, Mann-Whitney U test) (**C**) Comparison of elastic moduli (G’) between the control and nocodazole treated cells across the mid cytoplasmic region at 1 Hz (i) and 10 Hz (ii) frequencies. Comparison of dynamic viscosity (η) values between the control and nocodazole treated cells across the mid cytoplasmic region at 1 Hz (iii) and 10 Hz (iv) frequencies. (Nbeads > 60, Ncells > 10; ***p<0.001, Mann-Whitney U test) (**D**) Fractal dimension (FD) values of actin, microtubule (MT) and vimentin filaments in control and nocodazole treated cells across the apicobasal axis. (Mean ± SD, Ncells > 100; ***p<0.001, NS, non-significant, two sample T-test) (**E**) Representative confocal images of F-actin, microtubules (MT) and vimentin filaments in the control and nocodazole treated cells across the apicobasal axis. (Scale bar, 5 μm)

### Localised vimentin assembly stiffens mid cytoplasm in cytochalasin D treated cells

The disruption of F-actin resulted in the rise of cytoplasmic stiffness in the mid cytoplasmic region, with a concurrent increase in the density of vimentin filaments (**Figure 2A-i** and **Figure 2D-ii**). Therefore, we hypothesised that the elevated levels of vimentin filaments were responsible for the stiffening of the mid cytoplasm. To test this hypothesis, we knocked out vimentin using CRISPR-Cas9 and investigated the effect of Cyto D treatment on the mechanical properties of the mid cytoplasm in vimentin knockout (Vim^-/-^) cells. **Figure S4-A** validates the knockdown of vimentin. The Vim^-/-^ cells maintained the viscoelastic polarity of the cytoplasm along the apicobasal axis (**Figure 4A-i**). A comparison of **Figure 4A-i** and **Figure 1D** demonstrates that the polarity of compliance along the apicobasal axis slightly decreased at 1 Hz and increased at 10 Hz due to the reduction of vimentin filaments (**Figure S4-B**). The knockout of vimentin filaments made the basal and mid cytoplasm more compliant than the control cells (**Figure 4A-ii, iii**). Further, we observed a larger softening of the basal cytoplasm (75%) than the mid-cytoplasm (52%) at 1Hz due to vimentin knockout (**Figure S4-C**). However, at 10Hz, mid-cytoplasm had a larger softening than the basal cytoplasm (**Figure S4-C**). Cyto D treatment of Vim^-/-^ cells increased the compliance of the basal cytoplasm by an additional 5-fold at 1 Hz and 4-fold at 10 Hz (**Figure S4-D**). In contrast to the behaviour observed in wild-type cells, Vim^-/-^ cells did not exhibit reduced compliance or increased stiffness in the mid-cytoplasm when subjected to Cyto D treatment. This established the role of vimentin in the stiffening of the mid-cytoplasm upon Cyto D treatment as depicted in **Figure 2C**.

**Figure 4:**
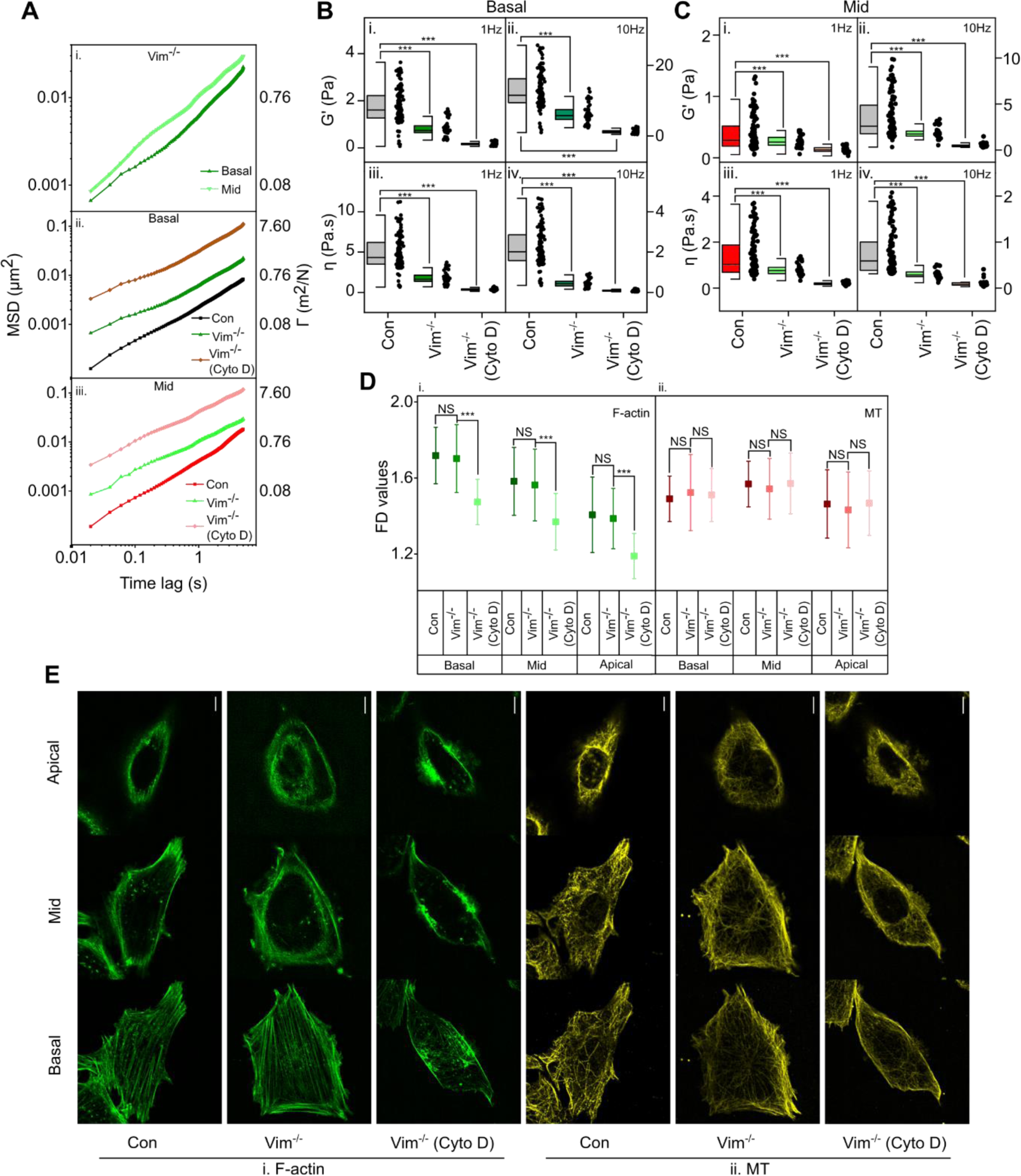
Localised vimentin assembly stiffens mid cytoplasm in cytochalasin D treated cells. (**A**) Mean MSD (left axis) and compliance, Γ (right axis) of the basal and mid cytoplasmic regions of Vim^-/-^ cells (i), comparison of Mean MSDs (left axis) and compliance, Γ (right axis) between the control, Vim^-/-^ and Cyto D treated Vim^-/-^ cells across the basal (ii) and mid (iii) cytoplasmic regions. (N_beads_ > 50, N_cells_ > 10) (**B**) Comparison of elastic moduli (G’) between the control, Vim^-/-^ and Cyto D treated Vim^-/-^ cells across the basal cytoplasmic region at 1 Hz (i) and 10 Hz (ii) frequencies. Comparison of dynamic viscosity (η) values between the control, Vim^-/-^ and Cyto D treated Vim^-/-^ cells across the basal cytoplasmic region at 1 Hz (iii) and 10 Hz (iv) frequencies. (N_beads_ > 50, N_cells_ > 10; ***p<0.001, Mann-Whitney U test) (**C**) Comparison of elastic moduli (G’) between the control, Vim^-/-^ and Cyto D treated Vim^-/-^ cells across the mid cytoplasmic region at 1 Hz (i) and 10 Hz (ii) frequencies. Comparison of dynamic viscosity (η) values between the control, Vim^-/-^ and Cyto D treated Vim^-/-^ cells across the mid cytoplasmic region at 1 Hz (iii) and 10 Hz (iv) frequencies. (N_beads_ > 50, N_cells_ > 10; ***p<0.001, Mann-Whitney U test) (**D**) Fractal dimension (FD) values of actin and microtubule (MT) filaments in control, Vim^-/-^ and Cyto D treated Vim^-/-^ cells across the apicobasal axis. (Mean ± SD, N_cells_ > 80; ***p<0.001, NS, non-significant, two sample T-test) (**E**) Representative confocal images of F-actin and microtubules (MT) filaments in the control, Vim^-/-^ and Cyto D treated Vim^-/-^ cells across the apicobasal axis. (Scale bar, 5 μm)

As anticipated, vimentin knockout caused a larger G’ reduction in the basal (55%) cytoplasm than in the mid (11%) cytoplasm (**Figure 4B**, **Figure 4C** and **Figure S4-E**). The knockout of vimentin increased the polarity of G’ and η along the apicobasal axis (**Figure S4-F**). Additionally, vimentin depletion increased the fluidity of the mid-cytoplasm at 1Hz (**Figure S4-G**). At 1 Hz, Cyto D treatment to Vim^-/-^ cells reduced the loss tangent (tan δ) of both the basal and mid cytoplasm. However, at 10 Hz, Cyto D treatment increased the tan δ of the basal cytoplasm and had a negligible impact on the tan δ of the mid cytoplasm. (**Figure S4-H**). Next, we investigated whether the knockout of vimentin affected the spatial arrangement of actin and microtubule filaments. The depletion of vimentin had an insignificant effect on the fractal dimension (FD) of the actin filament and microtubule networks (**Figure 4D**). **Figure 4E** compares the immunofluorescence images of the F-actin and microtubules at the basal, mid, and apical cytoplasm in wild-type (WT) cells with Vim^-/-^ cells. We observed no noticeable changes in the spatial organisation of actin and microtubule filaments due to vimentin knockout. The FD analysis (**Figure 4D**) supported this observation, suggesting that the knockout of vimentin had an insignificant effect on actin and microtubule filament networks in both the basal and mid-cytoplasm. Therefore, the altered mechanical properties of Vim^-/-^ cells can be attributed to the lack of vimentin filaments.

### Plectin directs the localised assembly of vimentin filaments in the mid-cytoplasm of cytochalasin D treated cells

The localised increase of vimentin filaments in Cyto D treated cells (**Figure 2D-iii** and **2E-iii**) suggested that the formation of new filaments is controlled spatially within the cytoplasm. To gain mechanistic insight into the localised growth of vimentin filaments in the mid-cytoplasm, we investigated whether the existing vimentin molecules within the filaments spatially reorganize or if the overexpression of vimentin in Cyto D-treated cells forms new filaments localised in the mid-cytoplasm. While the levels of actin (**Figure 5A**) and tubulin (**Figure 5B**) proteins were unaffected by Cyto D treatment, the expression levels of vimentin showed a significant increase (**Figure 5C**). Additionally, we observed a significant increase in the levels of vimentin mRNA due to Cyto D treatment (**Figure 5D**). This indicates that the disassembly of actin by Cyto D triggered the synthesis of additional vimentin proteins. Further, the newly formed vimentin molecules preferentially assembled into filaments in the mid cytoplasm. We speculated that crosslinkers such as plectin, which enable mechanosensitive crosslinking of vimentin with F-actin (Marks et al., 2022) and microtubules (Svitkina et al., 1996) in cells may control the localised growth of new vimentin filaments. To validate our assumption, we depleted plectin using plectin siRNA (**Figure 5E**) and investigated the effect of Cyto D treatment on the assembly of vimentin filaments in transiently plectin-depleted cells. Treatment of Cyto D increased the levels of vimentin protein (**Figure 5F**) and the mRNA (**Figure 5G**) in plectin-depleted cells. Unlike control cells, the treatment of Cyto D led to an increase of vimentin density in both the basal and mid cytoplasmic regions of plectin deficient cells (**Figure 5I and S5-D**). This established the role of plectin in facilitating the localised assembly of vimentin in the mid-cytoplasm after Cyto D treatment. Interestingly, we observed an increase in the density of actin filaments in the form of stress fibers in the basal cytoplasm of plectin- deficient cells (**Figure 5H and S5-D**). However, there was no observable change in the total protein level of actin in plectin-depleted cells (**Figure S5-A**). This suggests that the increase in actin filaments in the basal cytoplasm of plectin-depleted cells was orchestrated by the spatial reorganisation of filamentous actin rather than an overall increase in the level of total actin. Hence, we conclude that plectin helps maintain the relative abundance of actin filaments and is essential for the localised increase in the density of vimentin filaments in the mid-cytoplasmic region of Cyto D-treated cells. The reorganisation of actin and vimentin filaments in plectin-depleted cells resulted in a more gel-like consistency of the basal and mid cytoplasm at higher frequencies (**Figure S5-B** and **S5-C**).

**Figure 5:**
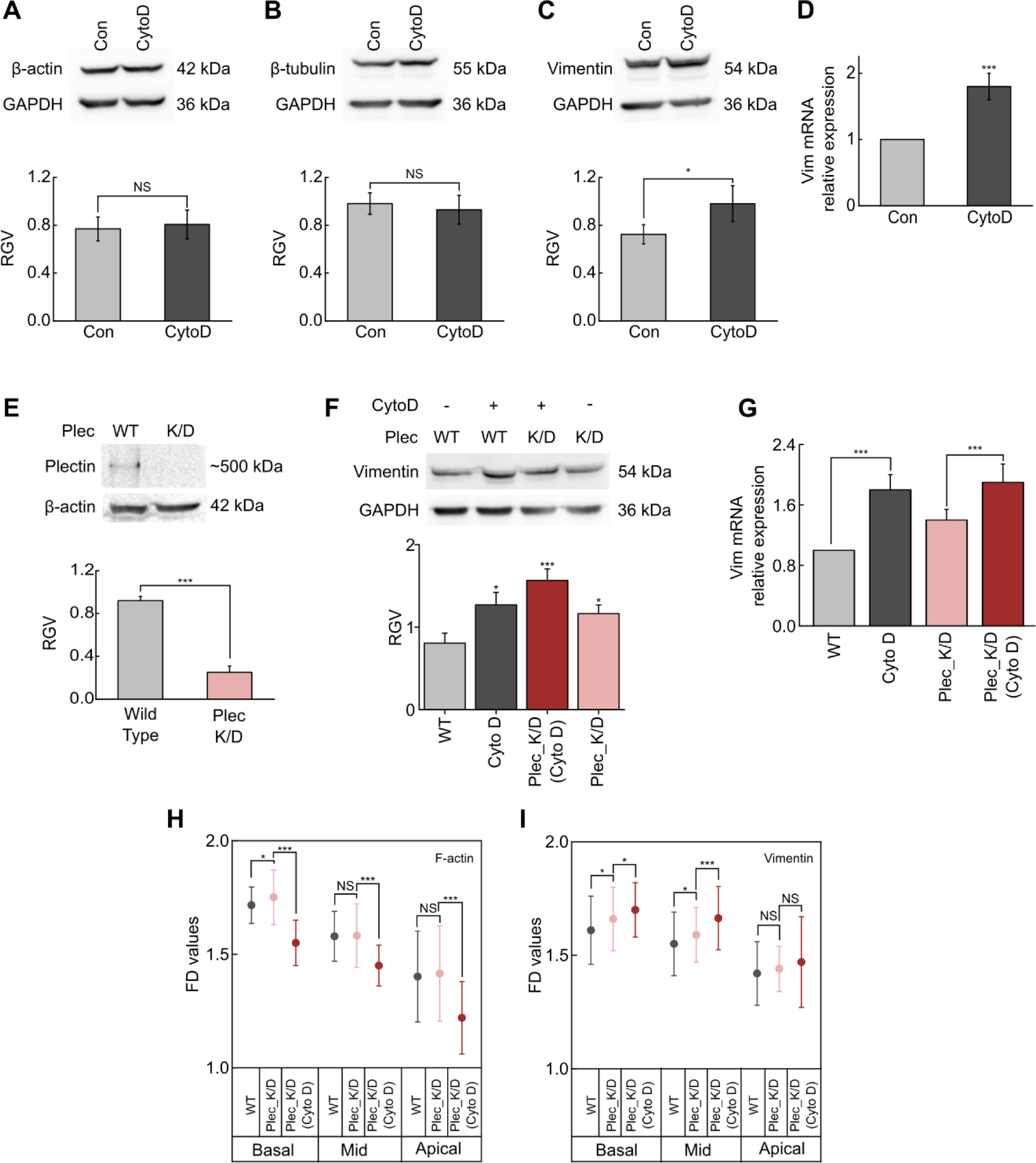
Plectin directs the localised assembly of vimentin filaments in the mid-cytoplasm of cytochalasin D treated cells. (**A**) Western blot analysis for β-actin protein levels post Cyto D (2μM, 1hr) treatment. (n=3, mean ± SD, NS, non-significant, two sample T test). (**B**) Western blot analysis for β-tubulin protein levels post Cyto D (2μM, 1hr) treatment. (n=3, mean ± SD, NS, non-significant, two sample T test). (**C**) Western blot analysis for vimentin protein levels post Cyto D (2μM, 1hr) treatment. (n=3, mean ± SD, *p<0.05, two sample T test). (**D**) Relative fold change of vimentin mRNA levels (as quantified by qPCR) post Cyto D treatment (2μM, 1hr). (n=3, mean ± SD, ***p<0.001, two sample T-test) (**E**) Western blotting analysis of plectin for confirmation of knockdown. (n=3, mean ± SD, ***p<0.001, two sample T test) (**F**) Western blot analysis of vimentin expression in wild type and plectin knockdown cells with and without Cyto D (2μM, 1hr) treatment. (n=3, mean ± SD, *p<0.05, ***p<0.001, two sample T test) (**G**) Relative fold change of vimentin mRNA levels (as quantified by qPCR) of wild type and plectin knockdown cells post Cyto D treatment (2μM, 1hr). (n=3, mean ± SD, *** p <0.001, two sample T-test). Fractal dimension (FD) values of actin (**H**) and vimentin (**I**) filaments in control, plectin depleted and Cyto D treated plectin depleted cells across the apicobasal axis. (Mean ± SD, N_cells_ > 80; *p<0.05, ***p<0.001, NS, non-significant, two sample T-test)

### Cyto D induced assembly of vimentin is dependent on ROS-mediated activation of NF-κΒ pathway

We observed that the disassembly of actin filaments by Cyto D caused an overexpression of vimentin via transcriptional upregulation (**Figure 5D**). The newly synthesized vimentin then assembled into filaments in the mid-cytoplasm (**Figure 2D-iii**). The treatment of Cyto D produces ROS (Kustermans et al., 2005), which is known to activate the transcription factor NF-κΒ (Morgan & Liu, 2011). Vimentin is a well reported target of NF-κΒ (Marsigliante et al., 2016; Zhou et al., 2019). Therefore, we speculated that Cyto D treatment activated NF-κΒ in NIH 3T3 cells through the generation of ROS, subsequently triggering the transcription of vimentin. Additionally, it remained unclear whether the elevation of ROS alone was sufficient or if the disassembly of actin filaments was also required to trigger the transcription and assembly of vimentin. Using the transient expression of EGFP-p65 in NIH 3T3 cells, we established that Cyto D treatment activated NF-κΒ within 15 minutes (**Figure 6A-i** and **6B-i**). We used an analogue of resveratrol, widely used as an NF-κB inhibitor, to establish that Cyto D activated NF-κB via the canonical pathway (**Figure 6A-ii**). Comparing the nuclear fraction of p65 in control cells with Cyto D-treated cells (**Figure 6C**) established that the Cyto D treatment activated the nuclear translocation of endogenous p65. Further, in the presence of the NF-κB inhibitor, Cyto D treatment failed to activate the transcription of vimentin (**Figure 6D-i**). This established that Cyto D activated the transcription of vimentin through NF-κB. The negligible change in the density of vimentin along the apicobasal axis post Cyto D treatment in cells pre-treated with the NF-κB inhibitor (**Figure 6E**, **Figure S6-B**) suggests that the increased filaments in the mid-cytoplasm arose from newly translated vimentin. Next, we explored whether Cyto D treatment led to an increase in the levels of intracellular ROS and further, whether the ROS was necessary to activate NF-κB. As expected, Cyto D treatment increased the intracellular ROS in NIH 3T3 cells (**Figure 6F**), which could be prevented by pre-treatment with vitamin C. Treatment of vitamin C alone neither altered the levels of intracellular ROS (**Figure 6F**) nor did it activate nuclear translocation of NF-κB (**Figure 6G-i and G-ii**). Vitamin C had a negligible effect on the density of vimentin filaments (**Figure S6-C**). Following that, we examined whether treatment of Cyto D in vitamin C pre-treated cells affected the transcription of vimentin. Despite the disassembly of actin filaments, the presence of vitamin C prevented Cyto D-mediated elevated expression of vimentin (**Figure 6H**). As expected, the presence of vitamin C prevented Cyto D-mediated increased assembly of vimentin filaments in the mid-cytoplasm (**Figure 6I**). This suggests that generation of ROS following the disruption of actin filaments is essential for the upregulation of vimentin in Cyto D treated cells. Interestingly, the elevation of ROS with H_2_O_2_ treatment led to a transcriptional (**Figure 6J**), as well as a translational (**Figure S6-G**) upregulation of vimentin (**Figure 6J**) and elevated density of the vimentin filaments across the apicobasal axis of cytoplasm (**Figure 6K and Figure S6-H**).

**Figure 6:**
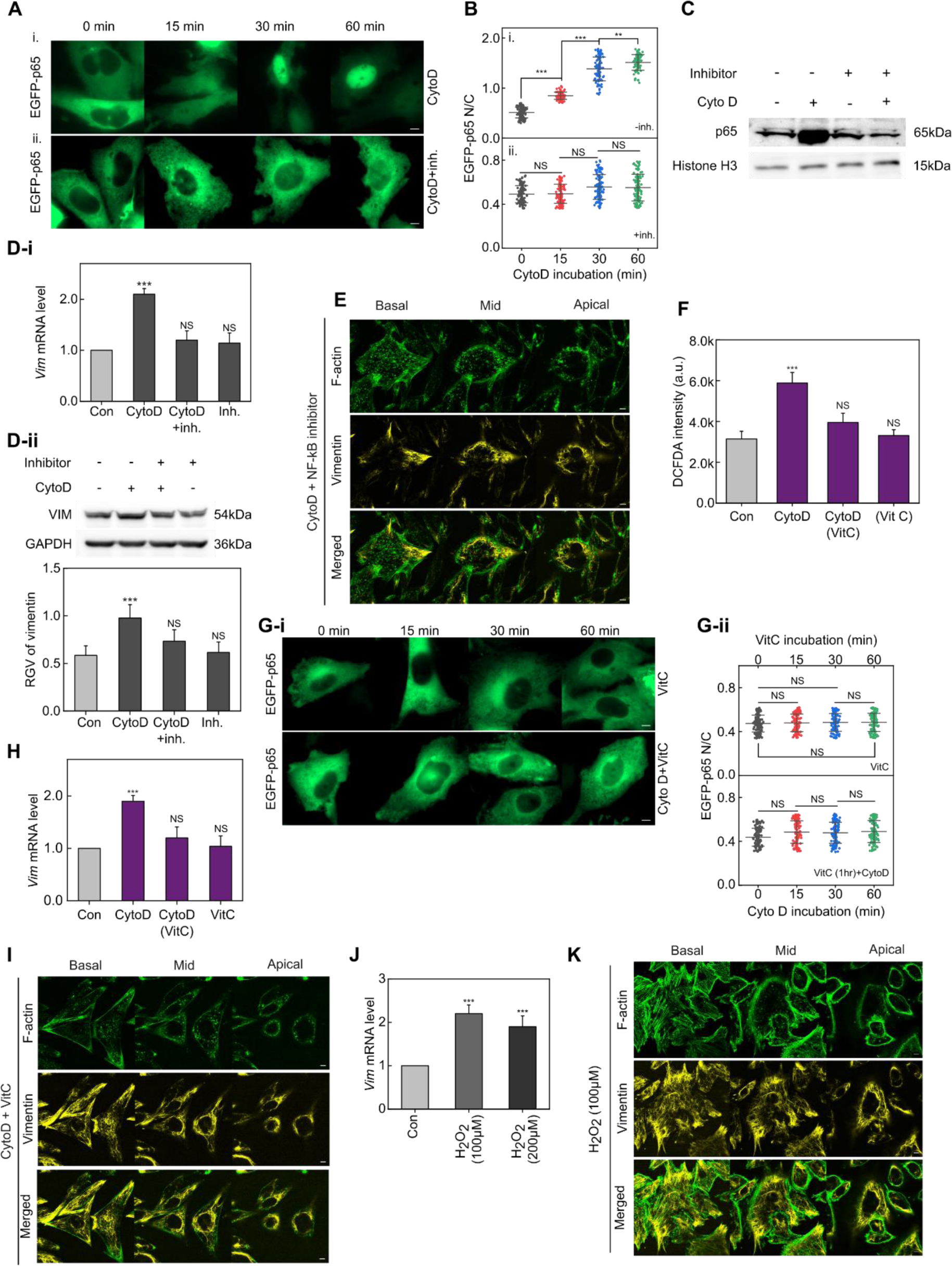
Cyto D induced assembly of vimentin is dependent on ROS-mediated activation of NF-κΒ pathway. (**A**) Representative confocal images of nuclear translocation of p65-GFP at different time points post Cyto D (2μM) treatment. (Scale bar, 5μm) (**B**) Nuclear to cytoplasmic (N/C) mean intensity ratio of p65 at different time points post Cyto D (2μM) treatment with and without NF-κΒ inhibitor. (N_Cells_ = 80-110 per condition, experimental replicates, n = 3, mean ± SD, **p<0.01, ***p<0.001, NS, non-significant two sample T test) (**C**) Western blotting analysis of the nuclear fraction of p65 after Cyto D (2μM, 1hr) treatment in the presence and absence of the NF-κΒ inhibitor. (**D-i**) Relative fold changes of vimentin mRNA levels (as quantified by qPCR) post-Cyto D (2μM, 1hr) treatment with and without NF-κΒ inhibitor pretreatment. (n = 3, mean ± SD, ***p<0.001, NS, non-significant, two sample T test) (**D-ii**) Western blot analysis for vimentin protein levels post-Cyto D (2μM, 1hr) treatment with and without NF-κΒ inhibitor pretreatment. (n = 3, mean ± SD, ***p<0.001, NS, non-significant, two sample T test) (**E**) Representative confocal images of F-actin and vimentin at the basal, mid and apical regions after Cyto D treatment (2μM, 1hr) in the presence of NF- κΒ inhibitor. (Scale bar, 5μm) (**F**) Mean DCFDA fluorescence levels of cells after Cyto D treatment (2μM, 1hr) pre-incubated with vitamin C (Vit C) (n = 3, mean ± SD, ***p<0.001, NS, non-significant, two sample T test). (**G-i**) Representative images of nuclear translocation of p65-GFP at different time spans of Vit C incubation (upper row) and p65-GFP nuclear translocation in cells after Cyto D (2μM) treatment for various durations following a 1-hour pre-treatment with Vit C (lower row). (Scale bar, 5μm). (**G-ii**) Nuclear to cytoplasmic (N/C) mean intensity ratio of p65 at different Vit C incubation periods (top) and N/C mean intensity ratio of p65 of cells after Cyto D (2μM) treatment for various durations following a 1hour pre-treatment with Vit C (bottom). (N_cells_ = 80-120 per condition, n = 3, mean ± SD, NS, non-significant, two sample T test). (**H**) Relative fold change of vimentin mRNA levels (as quantified by qPCR) post Cyto D treatment (2μM, 1hr) with and without Vit C pre-incubation. (n=3, mean ± SD, ***p<0.001, NS, non-significant, two sample T test). (**I**) Representative confocal images of F-actin and vimentin at the basal, mid and apical regions after Cyto D treatment (2μM, 1hr) in the presence of Vit C. (Scale bar, 5μm) (**J**) Relative fold change of vimentin mRNA levels (as quantified by qPCR) post H_2_O_2_ incubation (100μM and 200μM). (n=3, mean ± SD, ***p<0.001, two sample T test) (**K**) Representative confocal images of F-actin and vimentin at the basal, mid and apical regions after H_2_O_2_ incubation (100μM). (Scale bar, 5μm)

## Discussion

The components of the eukaryotic cytoskeleton, including actin, microtubules, and intermediate filaments, are interconnected, forming a complex meshwork of biopolymers (Hohmann & Dehghani, 2019). This cytoskeletal meshwork is one of the major regulators of the cytoplasmic mechanics (Fletcher & Mullins, 2010). The extent to which the assembly and distribution of different cytoskeletal components are interdependent, and their impact on the mechanical properties of the cytoplasm, is incompletely understood and requires further investigation. Vimentin can control the organisation of cortical actin in mitotic cells (Serres et al., 2020). Another study also emphasizes the importance of the interaction between vimentin filaments and cortical actin for normal cell division (Duarte et al., 2019). In our study, we observed that the organisation of actin and microtubule filaments in interphase fibroblasts remained unaffected by the knockout of vimentin (**Figure 4D** and **4E**). While the disassembly of actin filaments by Cyto D did not affect the density of microtubules, it resulted in an increased density of vimentin filaments in the mid-cytoplasm (**Figure 2D**), mediated by ROS-dependent activation of NF-κB. Cyto D treatment of cells pre-treated with either an NF-κB inhibitor (**Figure 6E-ii**) or a ROS quencher (**Figure 6I-ii**) prevented the increase in vimentin filaments in the mid-cytoplasm. This suggests that the disassembly of actin filaments alone does not trigger the assembly of vimentin filaments and requires ROS as an essential component for the activation of NF-κΒ. Nocodazole-induced disruption of microtubules did not affect the density of F-actin (**Figure 3D** and **3E-i**). However, intriguingly, nocodazole treatment reduced the density of vimentin filaments along the apicobasal axis (**Figure 3D** and **3E-iii**). Several reports suggest the importance of microtubules in the proper assembly and dynamics of vimentin filaments (Hookway et al., 2015; Prahlad et al., 1998). Therefore, we conclude that nocodazole-induced microtubule disruption impairs the proper assembly of vimentin filaments in NIH 3T3 fibroblasts. In U2OS cells, depletion of vimentin leads to an increase in stress fibers in the basal cytoplasm, along with a mild increase in actin monomers (Jiu et al., 2017). In contrast, depletion of vimentin in NIH 3T3 fibroblasts had a negligible effect on the stress fibres (**Figure 4D** and **4E-i**). This suggests that the interdependence of the relative distribution of different cytoskeletal components is cell type-dependent. Disassembly of actin by Cyto D caused a localised increase in vimentin filaments in the mid-cytoplasm (**Figure 2D** and **2E-iii**). We show that the localised assembly of vimentin in Cyto D-treated cells is dependent on the cytoskeletal crosslinker protein plectin (**Figure 5I** and **S5-D**), suggesting that plectin regulates the relative abundance of different cytoskeletal components in NIH 3T3 fibroblasts.

The rheological properties of the cytoplasm, specifically its stiffness (G’) and fluidity (η), are crucial for understanding cellular mechanics. In this manuscript, we investigate the polarity of these rheological properties along the apicobasal axis. Cells situated within a physically anisotropic extracellular microenvironment may exhibit distinct behaviors in their basal and apical regions. By examining the polarity of cytoplasmic stiffness and viscosity, we gain insights into how the mechanical properties of the cytoplasm influence cellular sensing and respond to extracellular mechanical cues in a heterogeneous 2D microenvironment. Furthermore, polarity in cytoplasmic stiffness is expected to influence the organization and migration of cells within tissues. Previous research has shown that fibroblast cells exhibit variable cytoplasmic stiffness at the leading and trailing edges during migration (Ford & Rajagopalan, 2018). Cortical F-actin is reported to enhance the stiffness of the cell cortex (Patteson et al., 2019). The distribution of actin, vimentin, and microtubules in different regions of the cytoplasm is inhomogeneous at macroscopic scales (**Figure 1H**). Thus, we speculate that the rheological properties of different cytoplasmic regions are differentially affected by various cytoskeletal filaments. Our study demonstrates that the anisotropic organisation of these cytoskeletal filaments results in polarised cytoplasmic stiffness and viscosity along the apicobasal axis in fibroblast cells cultured on glass substrate. The significantly higher relative abundance of actin and vimentin filaments in the basal cytoplasm (**Figure 1H**) suggests that these filaments likely govern the rheological properties of the basal cytoplasm. Cyto D treatment reduced the stiffness (**Figure 2B**) and increased the fluidity of the basal cytoplasm (**Figure S2A**) without altering the abundance of microtubules and vimentin (**Figure 2D**). Interestingly, depletion of vimentin (**Figure S4-A**) did not affect the abundance of actin or microtubule filaments in the basal cytoplasm (**Figure 4D**). Additionally, treatment with nocodazole, which disrupted both vimentin and microtubules in the basal cytoplasm (**Figure 3D**), significantly decreased the elastic modulus, G’ of the basal cytoplasm. This confirms that actin and vimentin filaments jointly dictate the rheological properties of the basal cytoplasm.

Longer actin filaments are absent in the mid-cytoplasm (**Figure 1G-i**). While we anticipate a substantial decrease in the density of actin filaments in the mid-cytoplasm compared to the basal cytoplasm in control cells, the FD of actin in the mid-cytoplasm (1.58) shows only a slight reduction compared to that in the basal cytoplasm (1.72). This smaller decrease in FD (**Figure 1H**) can be attributed to the presence of denser actin filaments in the cell cortex. Consequently, the FD values do not accurately reflect the actual density of F-actin in the perinuclear mid-cytoplasmic region. Therefore, we expect the rheological properties of the mid-cytoplasm to be less dependent on actin filaments and more influenced by vimentin and microtubules due to the absence of longer actin filaments. However, in Vim^-/-^ cells, Cyto D treatment altered the rheological properties of the mid-cytoplasm without affecting microtubules (**Figure 4B** and **4C**). We speculate that these changes in the rheological properties of the mid-cytoplasm in Vim^-/-^ cells following Cyto D treatment are due to alterations in the packing fraction of organelles (Gurel et al., 2014).

In cytoplasm where molecular motion of the cytoskeletal network is constrained by strong crosslinking and high filler content, a decrease in loss tangent with increasing frequency is expected, indicating reduced energy dissipation at higher frequencies. In contrast, an increase in loss tangent with frequency suggests a complex, time-dependent relaxation process, where energy dissipation mechanisms become more active at shorter time scales (higher frequencies). We infer that this behavior reflects the entanglement and weak crosslinking of cytoskeletal filaments. Since nocodazole treatment results in the loss of microtubules and a significant reduction in vimentin filaments while maintaining the density of actin filaments, we assume that the mechanical properties of the basal cytoplasm in nocodazole-treated cells are primarily governed by the actin filament network. The loss tangent of the basal cytoplasm in these cells resembles that of control cells, supporting our assumption. However, the loss tangent exhibits significantly different characteristics in the mid-cytoplasm (**Figure S3-C**). Therefore, we speculate that in nocodazole-treated cells, the mechanical properties of the mid-cytoplasm are not predominantly determined by cytoskeletal filaments but rather by the distribution of organelles. The comparable increase in compliance of the basal cytoplasm in control cells, as well as in vim^-/-^ cells following Cyto D treatment (**Figure S4-D**), suggests that vimentin and actin filaments independently govern the mechanical characteristics of the basal cytoplasm.

## Materials and methods

### Cell culture

NIH 3T3 fibroblast cells were obtained from NCCS, Pune, India. The cells were cultured in DMEM media (Himedia, #AL007G) supplemented with 10% FBS (Himedia, #RM10434) and 5% penicillin-streptomycin (Himedia, A001A) antibiotic cocktail solution. The cells were consistently maintained at 37°C and 5% CO_2_ inside a standard cell culture incubator. Throughout the experiments, the cells were continuously screened for mycoplasma contamination. For imaging purposes, cells were plated on confocal dishes coated with poly-L-lysine (Sigma, #P8920) for maximum adherence, ensuring that cell density never exceeded 60% confluency.

### Drug treatment

Cytochalasin D (Sigma, #C8273) was used at a concentration of 2 μM for 1 hour. Nocodazole (Merck, #487928) was used at a concentration of 10 μM for 1 hour. NF- κΒ inhibitor (CAY10512) was used at a concentration of 250 μM, 1 hour prior to the addition of cytochalasin D. Vitamin C (Merck, A92902) was used at a concentration of 200 μM for 1 hour before treating with cytochalasin D. Cells were incubated with two different concentrations (100 and 200 μM) of hydrogen peroxide (Merck, 107209) solution for 15 minutes. Thereafter, the H_2_O_2_-containing media was replaced with DMEM and the cells were further incubated for 40-60 minutes before carrying out experiments.

### Transfection

The EGFP-p65 plasmid (Addgene plasmid #111190) was a gift from Johannes A. Schmid (Schmid et al., 2000). Transfection was performed with FuGENE HD (Promega, #E2311) transfection reagent according to the manufacturer’s guidelines.

Cells were kept in serum-free DMEM for 12 hours prior to transfection. On the day of transfection, the confluency of the cells was around 60-80%.

### Knockdown of plectin

Knockdown of plectin was performed using predesigned Dicer-substrate siRNA (DsiRNA) from IDT (# mm.Ri.Plec.13.1). IDT negative control DsiRNA (# 51-01-14-03) was used as a control siRNA. siRNAs were transfected into cells using FuGENE HD (Promega, #E2311) transfection reagent according to the manufacturer’s protocol. Cells were incubated in siRNA-added media for a minimum of 72 hours to ensure efficient depletion of plectin. The silencing of plectin was validated by western blot analysis.

### Knockout of vimentin using CRISPR-Cas9

The vimentin knockout (Vim^-/-^) NIH 3T3 cell line was generated using the CRISPR- Cas9 technique. The guidelines for performing CRISPR-Cas9 mediated knockout were adopted from the work published by Ran et al., 2013.(Ran et al., 2013). The guide RNAs for vimentin were designed using the web-based CHOPCHOP tool (Montague et al., 2014). The following oligos were used to construct the dual-stranded guide RNA (gRNA) for vimentin:

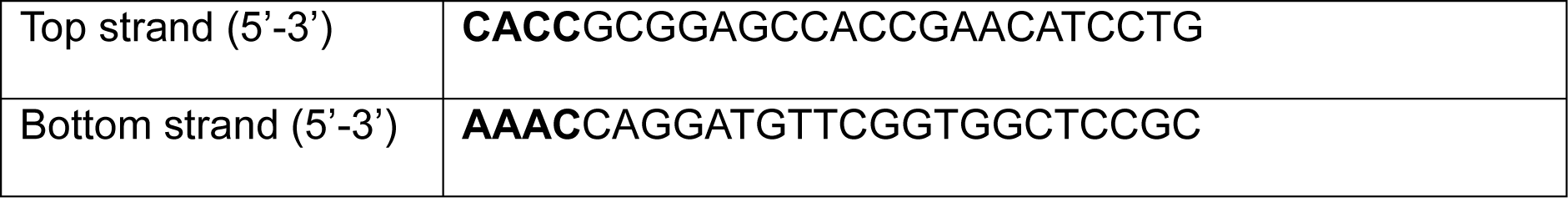

The bold letters denote the 5’ overhangs required for ligation into the pSpCas9(BB)- 2A-Puro V2.0 vector plasmid. The remaining 21 nucleotides of the top strand constitute the actual sequence of the gRNA: 5’-GCGGAGCCACCGAACATCCTG-3’.

The top and bottom strands were annealed together to create the dual-stranded gRNA, which was then cloned into the pSpCas9(BB)-2A-Puro V2.0 vector plasmid (Addgene ID #62988). The 5’ overhangs of these oligos were phosphorylated using T4 Polynucleotide Kinase (Thermo Scientific, #EK0031) to facilitate efficient ligation. The annealing of these two oligos was performed in a thermocycler, gradually reducing the temperature from 95°C to 25°C at a rate of 5°C per minute. Next, the gRNA construct was cloned into the pSpCas9(BB)-2A-Puro V2.0 vector for co-expression with Cas9. Prior to ligation, the pSpCas9(BB)-2A-Puro V2.0 vector was digested with BpiI restriction endonuclease enzyme (Thermo Scientific, #ER1011) at 37°C for 1 hour. There are two BpiI restriction sites in the pSpCas9(BB)-2A-Puro V2.0 vector. After digestion, a fragment between these two sites is excised, and the gRNA construct is inserted and ligated into the vector between these two sites. The ligation reaction was performed by incubating the gRNA insert and the pSpCas9(BB)-2A-Puro V2.0 vector at a ratio of 7:1 in the presence of T7 DNA ligase (New England Biolabs, # M0318S) enzyme at 25°C for 1 hour. After successful cloning of the gRNA into the pSpCas9(BB)-2A-Puro V2.0 vector, the entire construct was introduced into DH5α strain of competent bacteria using the standard bacterial transformation protocol. Following transformation, the bacteria were plated on Luria agar plates containing ampicillin. Bacteria which were successfully transformed with pSpCas9(BB)-2A-Puro V2.0 become resistant to ampicillin and gave rise to individual colonies. A few of these colonies were picked for colony PCR to confirm the presence of the cloned pSpCas9(BB)-2A-Puro V2.0 vector. Details of the primers used in the colony PCR is given below,

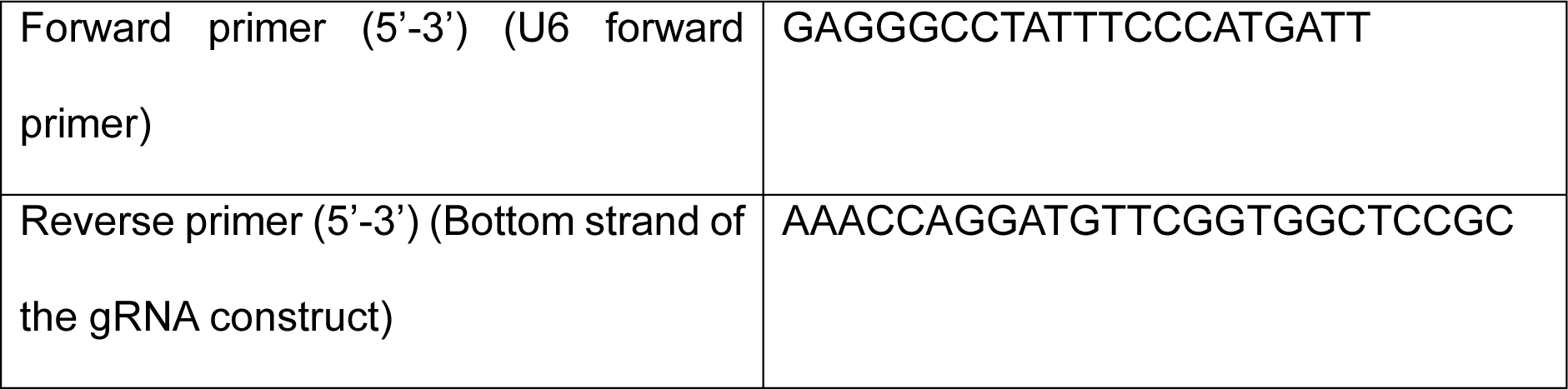

Following colony PCR, positive colonies were then plated onto another ampicillin- containing agar plate. One or two colonies from this plate were then picked up and grown in a liquid Luria broth culture medium at 37°C for overnight. The cloned gRNA + pSpCas9(BB)-2A-Puro V2.0 constructs were then isolated from the bacterial culture using a plasmid isolation kit (Macherey-Nagel, #740588.50). Following isolation, the cloned constructs were transfected into NIH 3T3 cells using FuGENE HD (Promega, #E2311) transfection reagent. pSpCas9(BB)-2A-Puro V2.0 plasmid contains puromycin resistance gene. Next, the transfected cells were subjected to puromycin (Sigma, #P8833) antibiotic treatment to isolate puromycin-resistant single-cell colonies. These colonies were then screened for the presence of vimentin using immunostaining and dot blot methods. Colonies with no vimentin expression were isolated and subsequently used in the later experiments.

### Western blotting

For Western blot experiments, protein lysate of each sample was prepared from approximately 3.2 × 10^6^ cells seeded on a 60 mm cell culture dish. Cells were washed three times with ice-cold PBS, then scraped and lysed in RIPA lysis buffer solution supplemented with 1X protease and phosphatase inhibitor cocktail (Merck, PPC1010). Protein concentrations were quantified using Bradford reagent. Blocking was performed using 5% BSA solution (1 hour at room temperature) and the washes were done using TBST buffer (Tris-buffered saline with 0.1% Tween 20). The antibodies were diluted in 5% BSA solution. The following primary antibodies were used: vimentin rabbit monoclonal antibody (dilution 1:1000; CST, #5741), β-tubulin rabbit polyclonal antibody (dilution 1:1000, Abclonal, #AC008), β-actin mouse monoclonal antibody (dilution 1:1000, SCBT, #sc-47778), GAPDH mouse monoclonal antibody (dilution 1:2000, Abclonal, #AC002), NF-κΒ p65 (L8F6) mouse monoclonal antibody (dilution 1:1000; CST, #6956), plectin mouse monoclonal antibody (dilution 1:500; SCBT, #sc- 33469), histone H3 rabbit polyclonal antibody (dilution 1:1000; Invitrogen, # PA5- 16183). The following horseradish peroxidase (HRP)-tagged secondary antibodies were used: anti-mouse secondary antibody (dilution 1:10000; Invitrogen, #31430) and anti-rabbit secondary antibody (dilution 1:10000; Invitrogen, #31460). Chemiluminescent signals from were developed by incubating the blots in a substrate solution containing luminol, coumaric acid, and H_2_O_2_, prepared in 1.5M Tris-HCl buffer (pH 8.8), for 30-40 seconds. The band intensities were quantified using ImageJ. The intensity ratios of the total protein to the loading control GAPDH were calculated.

### Protein isolation from nuclear and cytoplasmic fractions

Cells were harvested using a scraper after three washes with ice-cold PBS. The cell pellet was then resuspended in approximately 500 µl of cytoplasmic extraction buffer, supplemented with 0.05% NP-40 detergent. The samples were vortexed frequently to separate the nuclear fraction from the cytoplasmic fraction. Subsequently, the samples were centrifuged at 10,000 × g and 4°C for 15 minutes, and the supernatant, containing the cytoplasmic fraction, was collected. The remaining pellet was resuspended in 400 µl of nuclear extraction buffer and vortexed every few minutes for 40 minutes to lyse the nuclear membrane. Finally, the samples were centrifuged at 15,000 × g and 4°C for 15 minutes, and the supernatant, containing the nuclear protein fraction, was collected.

### Cell fixation and immunostaining

For immunostaining, cells were seeded on glass-bottom confocal dishes coated with either poly-L-lysine or fibronectin solution for optimal adhesion. The confluency of the seeded cells was kept below 70%. Following various drug treatments, cells were fixed in 4% PFA solution and permeabilized using 0.1% Triton X-100 in PBS. Blocking was performed using 2% BSA solution in PBS for 1 hour at room temperature. Cells were incubated with the following primary antibody solutions for 16 hours at 4°C: vimentin anti-rabbit monoclonal antibody (dilution 1:100; Cell Signalling Technology, #5741) and β-tubulin rabbit polyclonal antibody (dilution 1:200; Abclonal, #AC008). Alexa Fluor 488 and 594-tagged anti-rabbit secondary antibodies (CST) were used at a 1:500 dilution. F-actin was visualised using Alexa Fluor 488 conjugated phalloidin (dilution 1:200, Invitrogen, # A12379).

### qPCR analysis

The total RNA from NIH 3T3 cells was extracted using RNeasy mini kit (Qiagen, #74104). cDNA was synthesised from the RNA samples using Verso cDNA synthesis kit (Thermo Scientific, #AB1453A). qPCR was performed using Genious 2X SYBR Green Fast qPCR Mix (Abclonal, #RK21204) in an Applied Biosystems 7500 Fast Real-Time PCR system. The sequences of the primers used for the qPCR are provided in the table below,

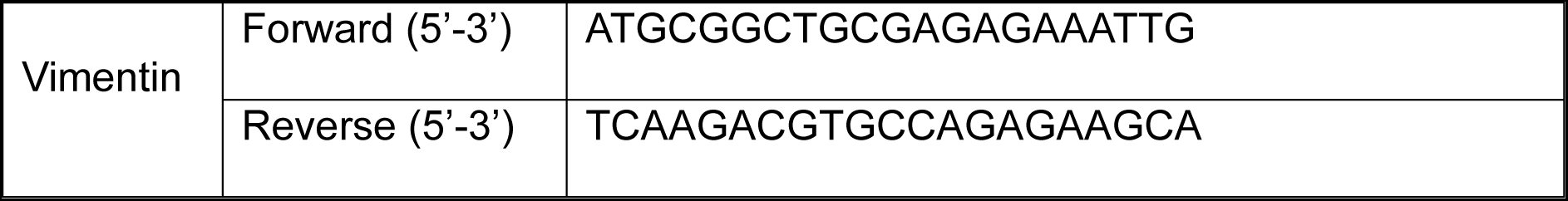
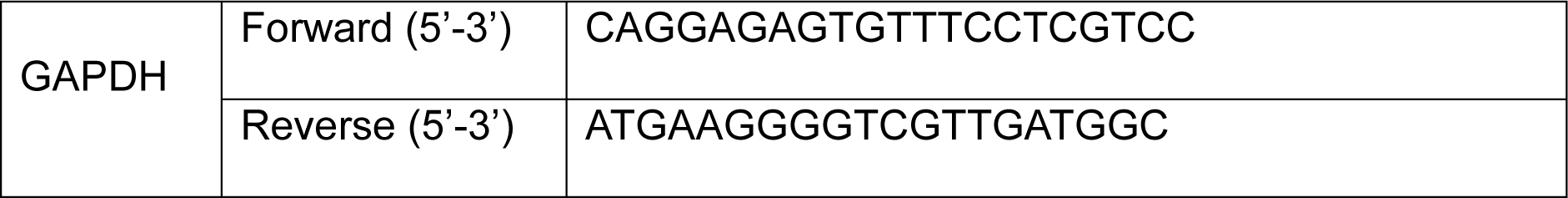

GAPDH was used as the housekeeping gene to normalise the expression levels of vimentin.

### Measurement of ROS

Intracellular ROS levels under various conditions were quantified using DCFDA (Merck) solution. Cells were seeded on a 96-well plate at a density of approximately 25000 cells per well and allowed to adhere properly for overnight. Thereafter, the cells in each well were incubated with 100μL of DCFDA (20 μM) solution prepared in serum- free DMEM media for 1 hour. Next, the DCFDA solution was washed out, and the cells were treated with the desired drug solutions (prepared in DMEM) for the appropriate duration. Subsequently, the fluorescence intensity of DCFDA was measured using a fluorescence plate reader (Ex/Em = 485/535 nm). The fluorescent intensity correlates with ROS levels.

### Microscopy and image analysis

Immunofluorescent images of actin, vimentin, and microtubule filaments were captured using a Zeiss LSM 880 confocal microscope with a 63X (NA 1.4) oil immersion objective (Zeiss, Thornwood, NY). Z-stack images were captured with an interval of 0.45 μm between each slice. To minimize cell-to-cell variation, cells with similar heights were selected, maintaining a consistent number of z-slices (8-10) for each cell. Fluorescent images of cells expressing EGFP-p65 were captured using a Zeiss AXIO Observer.Z1 wide-field epifluorescence microscope with a 63X oil immersion objective. Live cell imaging was performed on a heated microscope stage (+37°C) in presence of 5% CO_2_. Images were acquired using either Zen 2.0 or μManager software. Calculation of nuclear-to-cytoplasmic (N/C) intensity ratios of EGFP-p65 was carried out by selecting regions of interest (ROIs) from cytoplasmic and nuclear regions, and their mean intensity values were quantified using ImageJ.

### Fractal dimension analysis

Fractal dimension (FD) values of the actin, vimentin, and microtubule filaments were calculated using the FracLac (v2.5) plugin (Karperien, 1999) for ImageJ. 16-bit confocal images were converted into 8-bit binary images. Thresholding was performed using the Otsu method. During thresholding, efforts were made to ensure maximum visibility of cytoskeletal filaments while minimising background noise.

### Particle tracking microrheology (PTM)

The entire procedure of PTM was conducted as described in the standard protocol (Tseng et al., 2002; Wirtz, 2009). Fluorescently tagged carboxylate modified microspheres (Invitrogen, #F8811) having diameter of 200nm were ballistically shot inside the cell using a gene gun system (Bio-Rad, Helios® Gene gun system). After the shooting, the cells were washed with 1X PBS solution to wash away unbound beads and detached cells. The cells were then kept in a cell culture incubator under standard conditions (37°C, 5% CO_2_) for 16 hours in serum-free DMEM medium to allow healing from the injuries caused by the shooting of the microspheres. The cells containing fluorescent microspheres were subsequently positioned on the microscope stage at 37°C. The movements of the microspheres were recorded in the form of movies using a high-speed Hamamatsu Orca Flash 4.0 CMOS camera and μManager software (A. Edelstein et al., 2010; A. D. Edelstein et al., 2014). Each movie comprised 1000 frames, spanning a total duration of 20 seconds, with an exposure time of 20 milliseconds. The trajectories of the microspheres were extracted by tracking their movements using the MOSAICsuite plugin (Sbalzarini & Koumoutsakos, 2005) of ImageJ. At least 5-10 beads from each cell were tracked to obtain the rheological properties of the cytoplasm. The Mean Squared Displacement (MSD) and compliance (Γ) values were computed from these trajectories using a MATLAB script. MSDs having slopes greater than 1 were filtered out. While the movement of the microspheres was recorded for 1000 frames, only the first 250 frames were considered for calculating the MSDs to avoid potential errors from later data points in the trajectories. Subsequently, the complex shear modulus (G*(ω)), viscous modulus (G’’), and elastic modulus (G’) were derived from the MSDs using another MATLAB script.

## Acknowledgements

DR would like to thank Mahesh Agarwal and Parijat Biswas for their valuable insights into the principles and methodology of particle tracking microrheology. Special thanks to Mrs. Debapriya Ghatak for her assistance with the confocal microscopy.

## Author contributions

**DKS**: Conceptualisation, Funding acquisition, Resources, Supervision, Writing – original draft, Writing – reviewing and editing. **DR**: Conceptualisation, performed experiments, data analysis, data interpretation, wrote MATLAB scripts, Writing - reviewing and editing, preparation of figures.

## Supplementary Information

**Figure S1:**
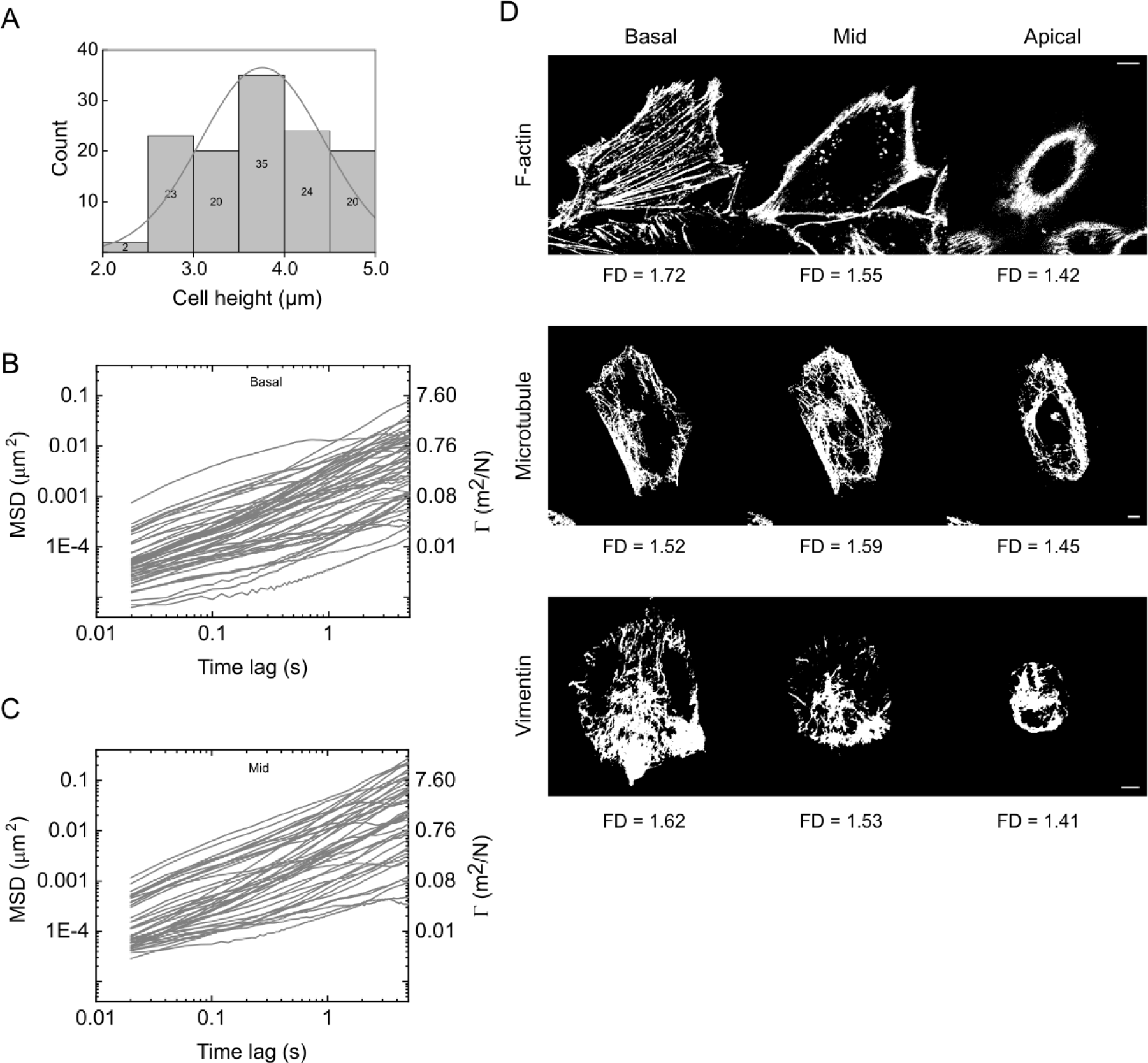
(A) Histogram profile of the height of NIH 3T3 cells (N_cells_ > 100). MSDs (left axis) and compliance, Γ (right axis) values of individual microspheres dispersed in the basal (B) and mid (C) cytoplasm of control NIH 3T3 cells. (N_beads_ > 60, N_cells_ > 10). (D) Representative binary images of actin, vimentin and microtubule filaments across the apicobasal axis of control NIH 3T3 cells and their corresponding fractal dimension (FD) values.

**Figure S2:**
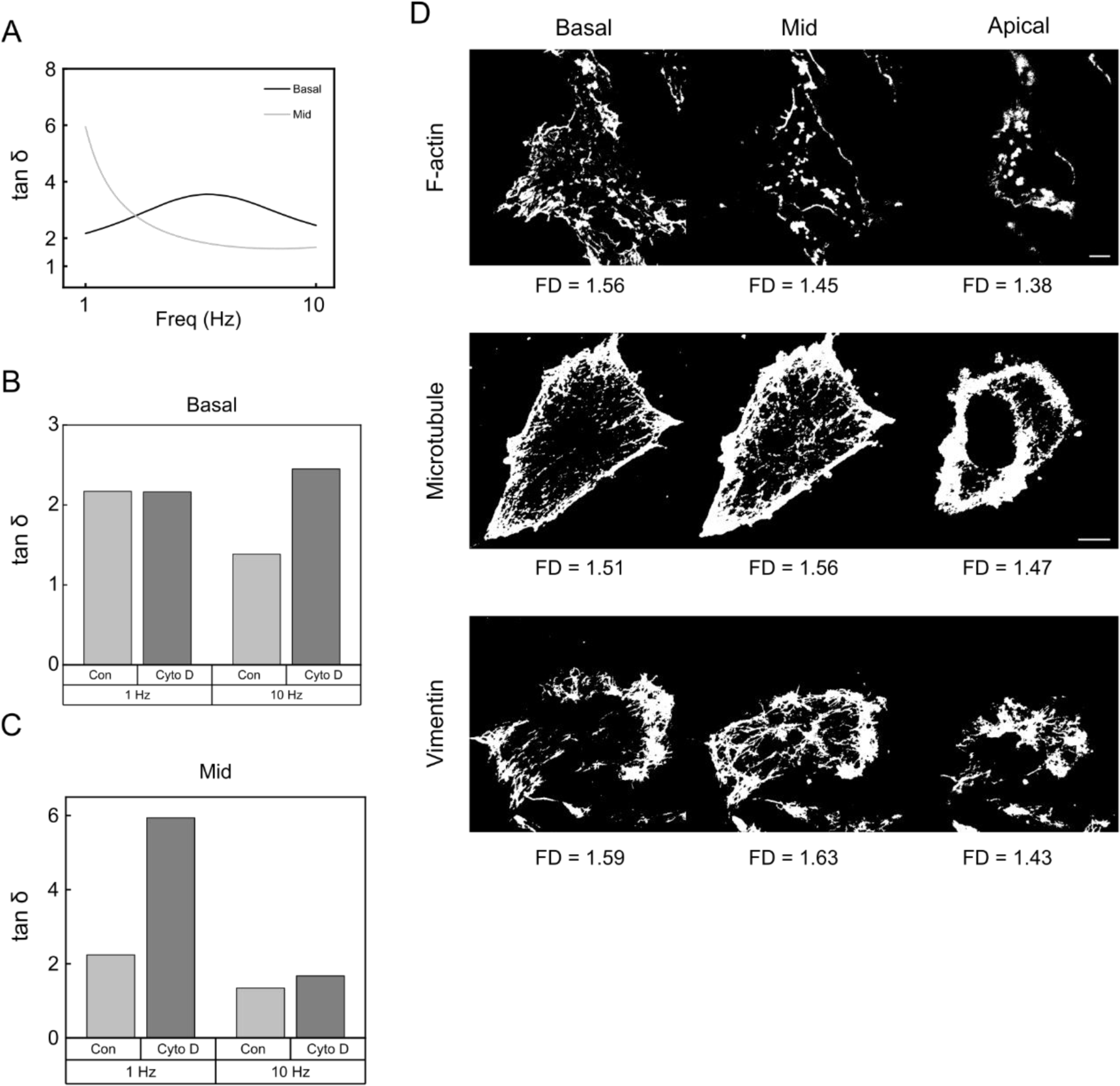
(**A**) Mean loss tangent values of the basal and mid cytoplasm of cyto D treated cells. (**B**) Comparison of tan δ values between control and cyto D treated cells at the basal cytoplasmic region at 1Hz and 10Hz. (**C**) Comparison of tan δ values between control and cyto D treated cells at the mid cytoplasmic region at 1Hz and 10Hz frequencies. (**D**) Representative binary images of actin, vimentin and microtubule filaments across the apicobasal axis of cyto D treated NIH 3T3 cells and their corresponding fractal dimension (FD) values.

**Figure S3:**
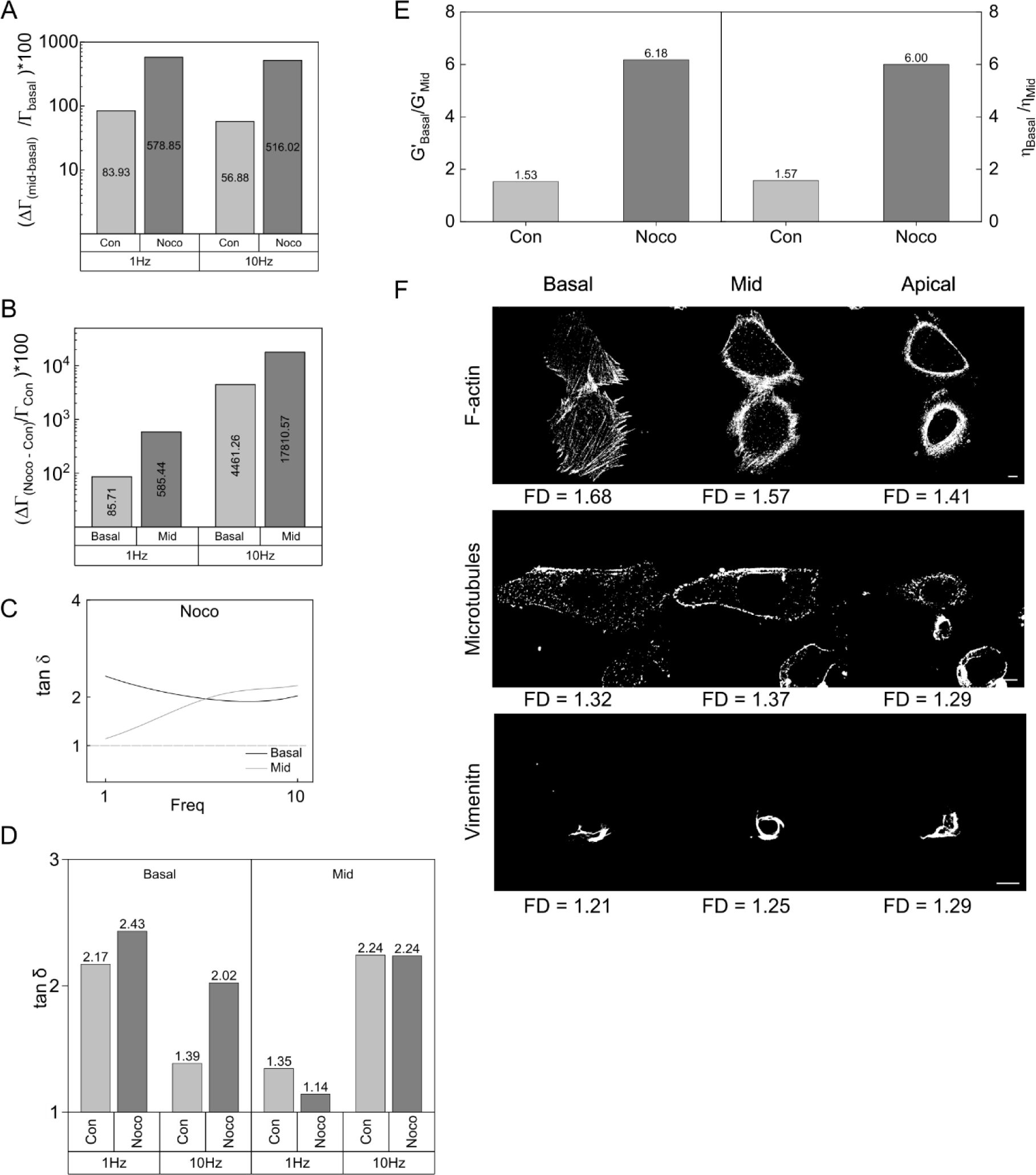
(**A**) Relative fold change of the compliance, Γ of the mid cytoplasm compared to the basal cytoplasm of control and nocodazole treated cells at 1Hz and 10Hz. (**B**) Relative fold change of the compliance, Γ of the nocodazole treated cells compared to the control cells in the basal and mid cytoplasm at 1Hz and 10Hz. (**C**) Loss tangent (tan δ) values of the basal and mid cytoplasmic regions of nocodazole treated cells. (**D**) Comparison of tan δ values between the control and nocodazole treated cells across the basal and mid cytoplasm at 1Hz and 10Hz. (**E**) Basal to mid cytoplasmic ratio of elastic modulus (G’) and dynamic viscosity (η) in control and nocodazole treated cells. (**F**) Representative binary images of actin, vimentin and microtubule filaments across the apicobasal axis of nocodazole treated NIH 3T3 cells and their corresponding fractal dimension (FD) values.

**Figure S4:**
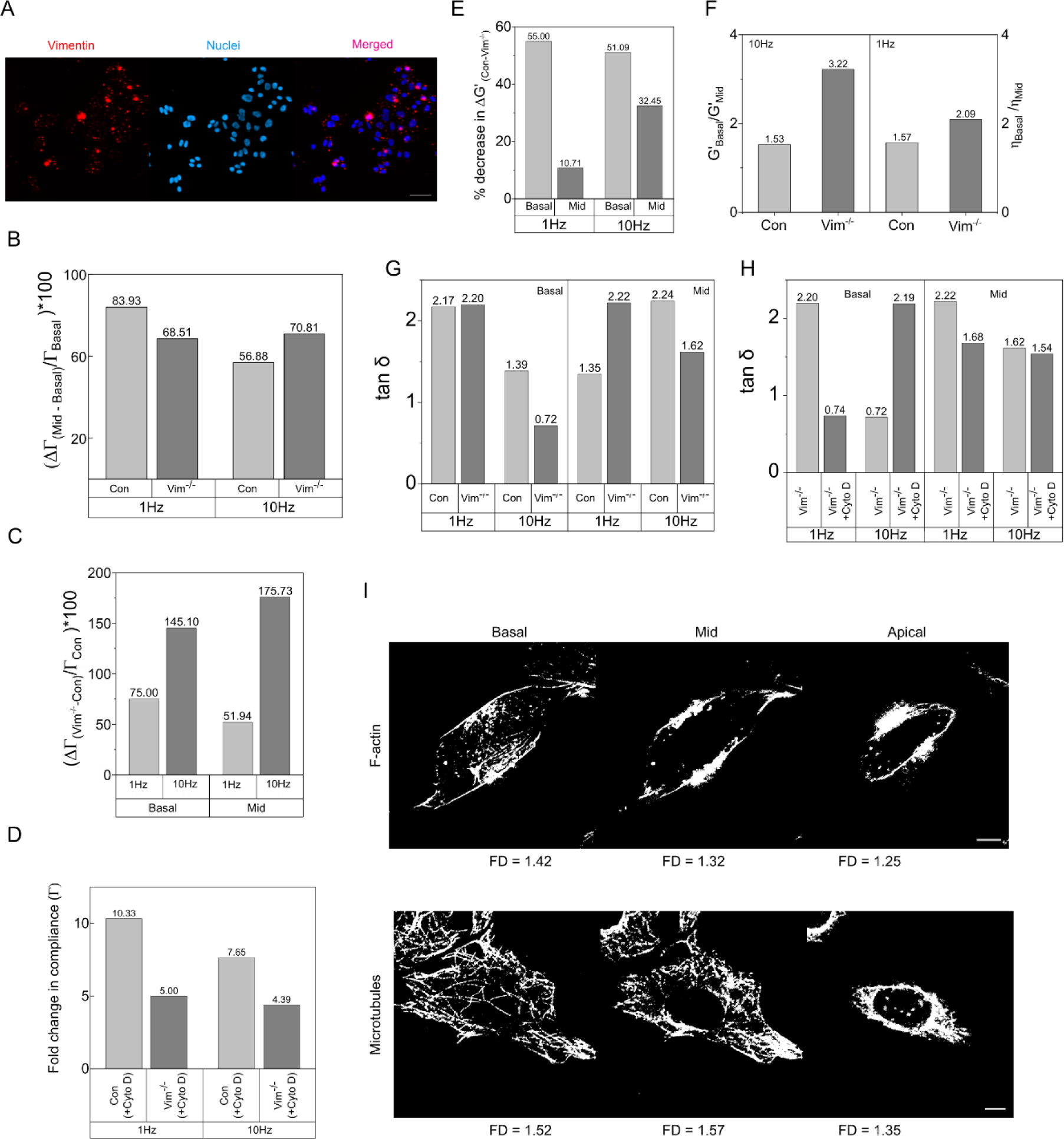
(**A**) Representative confocal image validating the knockout of vimentin from NIH 3T3. (Scale bar, 50μm) (**B**) Relative fold change of the compliance, Γ of the mid cytoplasm compared to the basal cytoplasm of control and Vim^-/-^ cells at 1Hz and 10Hz. (**C**) Relative fold change of the compliance, Γ of the Vim^-/-^ cells compared to the control cells in the basal and mid cytoplasm at 1Hz and 10Hz. (**D**) Fold change in the compliance, Γ of the basal region of control and Vim^-/-^ cells after cyto D treatment at 1Hz and 10Hz. (**E**) Percentage decrease in elastic modulus (G’) values upon knockout of vimentin across the basal and mid cytoplasm at 1Hz and 10Hz. (**F**) Basal to mid cytoplasmic ratio of elastic modulus (G’) and dynamic viscosity (η) in the control and Vim^-/-^ cells. (**G**) Comparison of tan δ values between the control and Vim^-/-^ cells across the basal and mid cytoplasm at 1Hz and 10Hz. (**H**) Comparison of tan δ values between the Vim^-/-^ and cyto D treated Vim^-/-^ cells across the basal and mid cytoplasm at 1Hz and 10Hz. (**I**) Representative binary images of actin and microtubule filaments across the apicobasal axis of cyto D treated Vim^-/-^ cells and their corresponding fractal dimension (FD) values.

**Figure S5:**
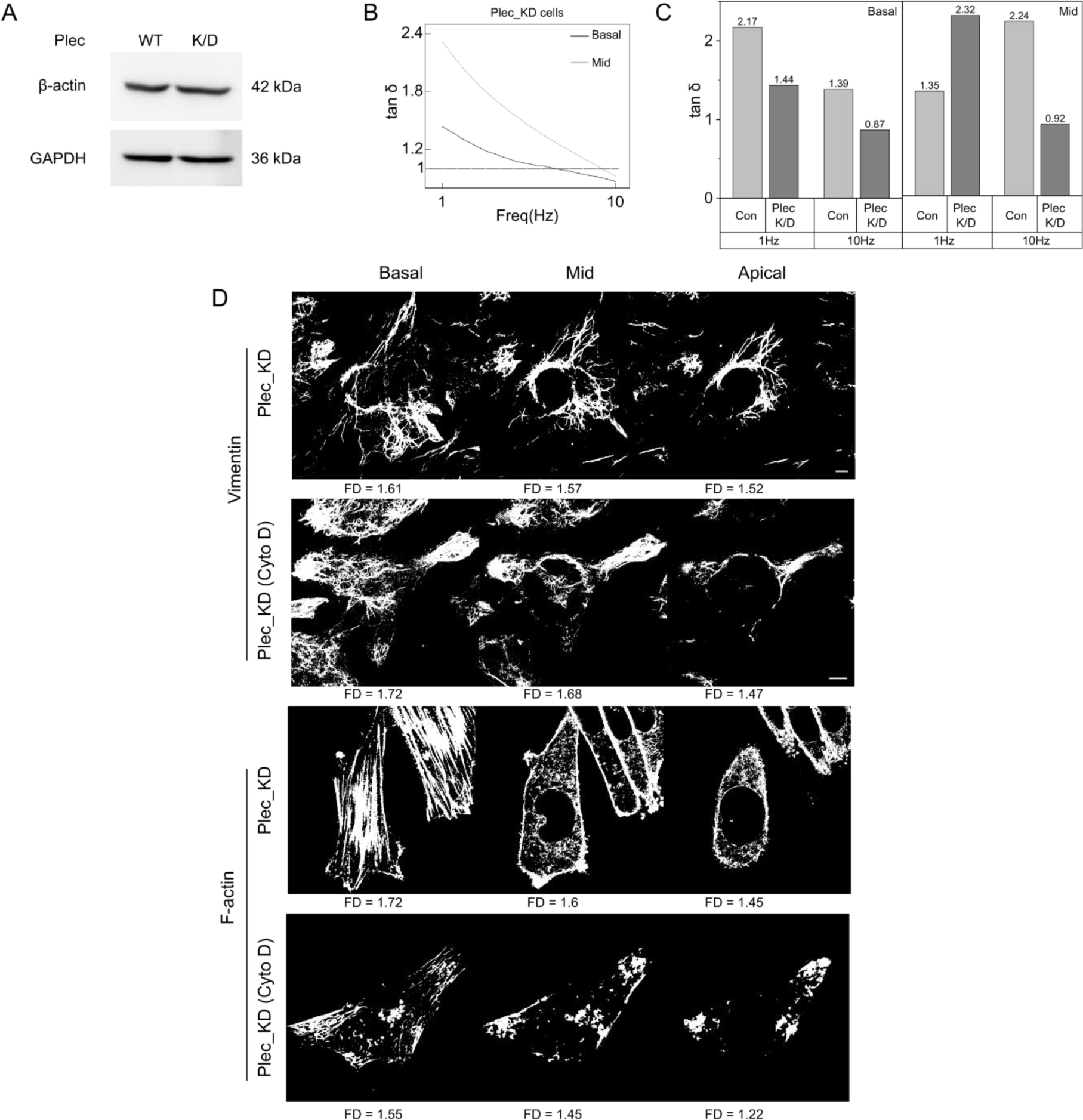
(**A**) Western blot analysis of beta-actin in control and plectin depleted NIH 3T3 cells. (**B**) Mean loss tangent values (tan δ) of the basal and mid cytoplasm of plectin depleted NIH 3T3 cells. (**C**) Comparison of tan δ values between the control and plectin depleted cells across the basal and mid cytoplasm at 1Hz and 10Hz. (**D**) Representative binary images of vimentin and actin filaments across the apicobasal axis of plectin-depleted and cyto D-treated plectin-depleted NIH 3T3 cells, along with their corresponding fractal dimension (FD) values.

**Figure S6:**
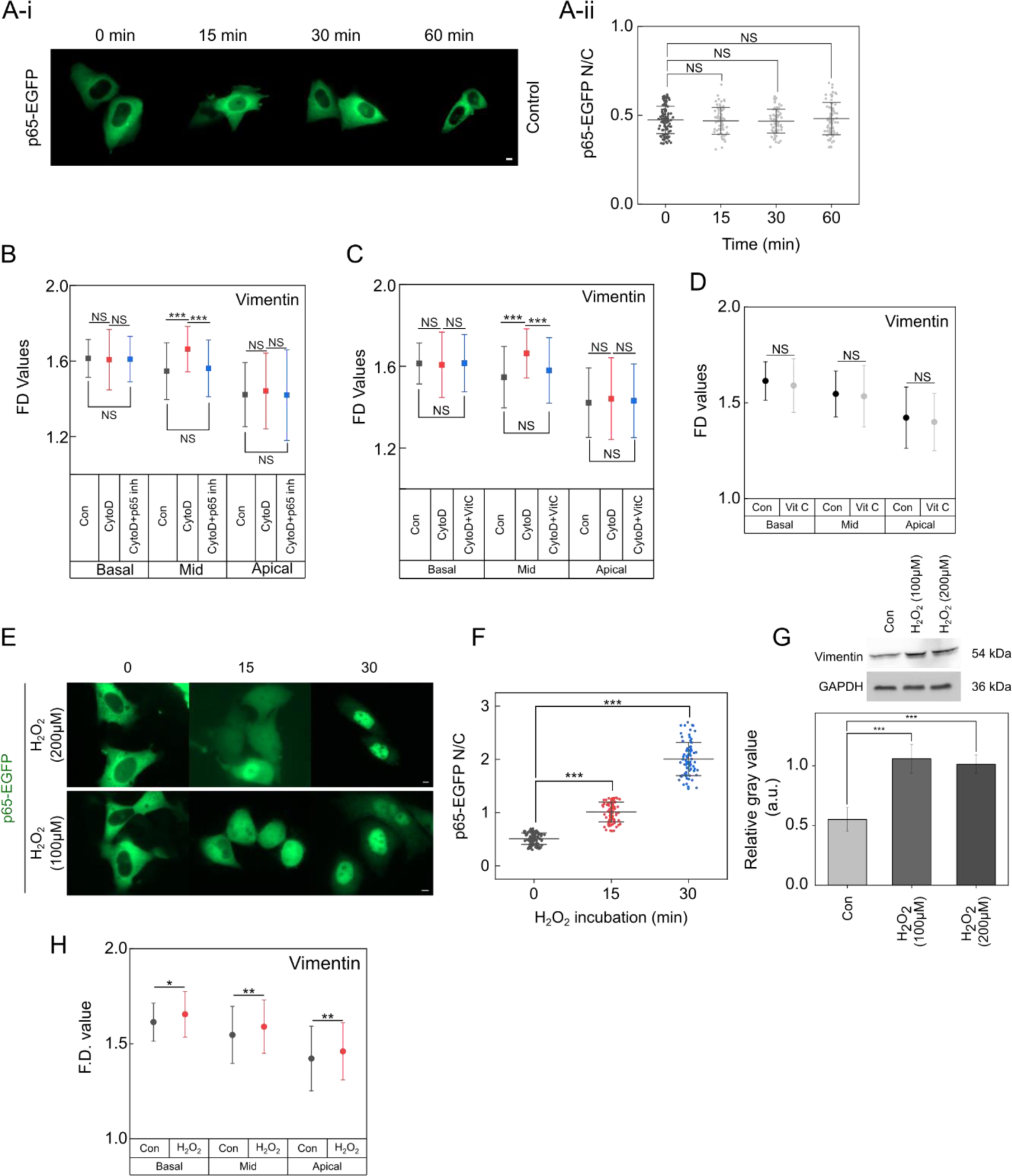
(**A**) Tracking of EGFP-p65 localization across different time points (i) (Scale bar, 5μm) and the corresponding nuclear to cytoplasmic (N/C) mean intensity ratio values (ii) (Mean ± SD, N_cells_ > 50, NS, non-significant, two sample T test). (**B**) Quantification of FD values of vimentin in control, cyto D treated and cyto D with p65 inhibitor pretreated cells across the basal, mid and apical cytoplasm. (N_cells_ = 100 -140 per condition, n = 3, mean ± SD, ***p<0.001, NS, non-significant, two sample T test) (**C**) Quantification of FD values of vimentin in control, cyto D treated and cyto D with vitamin C pretreated cells across the basal, mid and apical cytoplasm. (N_cells_ > 80, n = 3, mean ± SD, ***p<0.001, NS, non-significant, two sample T test) (**D**) FD values of control and vitamin C treated cells across the apicobasal axis. (N_cells_ > 100, n = 3, mean ± SD, NS, non-significant, two sample T test). (**E**) Nuclear translocation of EGFP-p65 at different time points following H_2_O_2_ incubation at two different concentrations. (Scale bar, 5μm) (**F**) Nuclear to cytoplasmic (N/C) mean intensity ratio of EGFP-p65 at different time points following H_2_O_2_ incubation. (N_cells_ > 60, ***p<0.001, two sample T test) (**G**) Western blot analysis of vimentin following H_2_O_2_ incubation at two different concentrations. (n = 3, mean ± SD, ***p<0.001, two sample T test) (**H**) FD analysis of vimentin in control and H_2_O_2_ treated cells across the apicobasal axis. (N_cells_ > 80, *p<0.05, **p<0.01, two sample T test)

**Figure S7:**
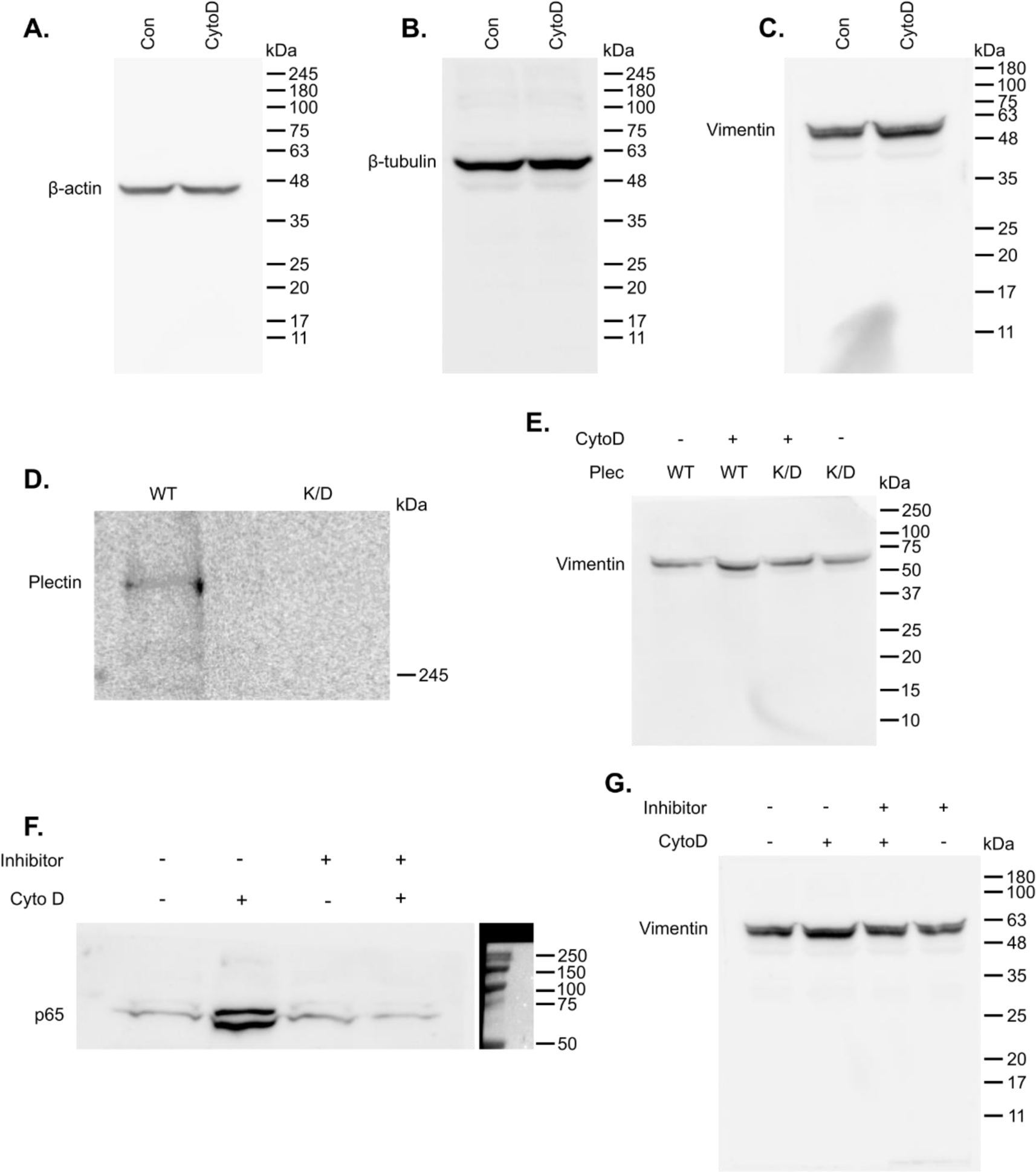
Blot transparency. (**A**) Complete western blot of β-actin represented in Figure 5A. (**B**) Complete western blot of β-tubulin represented in Figure 5B. (**C**) Complete western blot of vimentin represented in Figure 5C. (**D**) Complete western blot of plectin represented in Figure 5E. (**E**) Complete western blot of vimentin represented in Figure 5F. (**F**) Complete western blot of p65 represented in Figure 6C. (**G**) Complete western blot of vimentin represented in Figure 6D-ii.

## References

1. Bausch, A. R., Ziemann, F., Boulbitch, A. A., Jacobson, K., & Sackmann, E. (1998). Local measurements of viscoelastic parameters of adherent cell surfaces by magnetic bead microrheometry. Biophysical Journal, 75(4), 2038–2049. 10.1016/S0006-3495(98)77646-5

2. Bray, D. (2000). *Cell Movements: From Molecules to Motility* (2nd ed.). Garland Science. 10.4324/9780203833582

3. Brown, S. S., & Spudich, J. A. (1981). Mechanism of action of cytochalasin: Evidence that it binds to actin filament ends. The Journal of Cell Biology, 88(3), 487–491. 10.1083/jcb.88.3.487

4. Casella, J. F., Flanagan, M. D., & Lin, S. (1981). Cytochalasin D inhibits actin polymerization and induces depolymerization of actin filaments formed during platelet shape change. Nature, 293(5830), 302–305. 10.1038/293302a0

5. Cooper, J. A. (1987). Effects of cytochalasin and phalloidin on actin. The Journal of Cell Biology, 105(4), 1473–1478. 10.1083/jcb.105.4.1473

6. Duarte, S., Viedma-Poyatos, Á., Navarro-Carrasco, E., Martínez, A. E., Pajares, M. A., & Pérez-Sala, D. (2019). Vimentin filaments interact with the actin cortex in mitosis allowing normal cell division. Nature Communications, 10(1), 4200. 10.1038/s41467-019-12029-4

7. Edelstein, A., Amodaj, N., Hoover, K., Vale, R., & Stuurman, N. (2010). Computer Control of Microscopes Using µManager. Current Protocols in Molecular Biology, 92(1). 10.1002/0471142727.mb1420s92

8. Edelstein, A. D., Tsuchida, M. A., Amodaj, N., Pinkard, H., Vale, R. D., & Stuurman, N. (2014). Advanced methods of microscope control using μManager software. Journal of Biological Methods, 1(2), e10. 10.14440/jbm.2014.36

9. Ellis, R. J. (2001). Macromolecular crowding: Obvious but underappreciated. Trends in Biochemical Sciences, 26(10), 597–604. 10.1016/s0968-0004(01)01938-7

10. Fels, J., Orlov, S. N., & Grygorczyk, R. (2009). The hydrogel nature of mammalian cytoplasm contributes to osmosensing and extracellular pH sensing. Biophysical Journal, 96(10), 4276–4285. 10.1016/j.bpj.2009.02.038

11. Fletcher, D. A., & Mullins, R. D. (2010). Cell mechanics and the cytoskeleton. Nature, 463(7280), 485–492. 10.1038/nature08908

12. Ford, A. J., & Rajagopalan, P. (2018). Measuring Cytoplasmic Stiffness of Fibroblasts as a Function of Location and Substrate Rigidity Using Atomic Force Microscopy. ACS Biomaterials Science & Engineering, 4(12), 3974–3982. 10.1021/acsbiomaterials.8b01019

13. Guigas, G., Kalla, C., & Weiss, M. (2007). Probing the nanoscale viscoelasticity of intracellular fluids in living cells. Biophysical Journal, 93(1), 316–323. 10.1529/biophysj.106.099267

14. Gurel, P. S., Hatch, A. L., & Higgs, H. N. (2014). Connecting the Cytoskeleton to the Endoplasmic Reticulum and Golgi. Current Biology, 24(14), R660–R672. 10.1016/j.cub.2014.05.033

15. Hohmann, T., & Dehghani, F. (2019). The Cytoskeleton-A Complex Interacting Meshwork. Cells, 8(4), 362. 10.3390/cells8040362

16. Hookway, C., Ding, L., Davidson, M. W., Rappoport, J. Z., Danuser, G., & Gelfand, V. I. (2015). Microtubule-dependent transport and dynamics of vimentin intermediate filaments. Molecular Biology of the Cell, 26(9), 1675–1686. 10.1091/mbc.E14-09-1398

17. Hu, J., Jafari, S., Han, Y., Grodzinsky, A. J., Cai, S., & Guo, M. (2017). Size- and speed-dependent mechanical behavior in living mammalian cytoplasm. Proceedings of the National Academy of Sciences of the United States of America, 114(36), 9529–9534. 10.1073/pnas.1702488114

18. Jiu, Y., Peränen, J., Schaible, N., Cheng, F., Eriksson, J. E., Krishnan, R., & Lappalainen, P. (2017). Vimentin intermediate filaments control actin stress fiber assembly through GEF-H1 and RhoA. Journal of Cell Science, 130(5), 892–902. 10.1242/jcs.196881

19. Karperien, A. (1999). FracLac for ImageJ. https://imagej.net/ij/plugins/fraclac/FLHelp/Introduction.htm

20. Kustermans, G., El Benna, J., Piette, J., & Legrand-Poels, S. (2005). Perturbation of actin dynamics induces NF-kappaB activation in myelomonocytic cells through an NADPH oxidase-dependent pathway. The Biochemical Journal, 387(Pt 2), 531–540. 10.1042/BJ20041318

21. Liu, C.-Y., Lin, H.-H., Tang, M.-J., & Wang, Y.-K. (2015). Vimentin contributes to epithelial-mesenchymal transition cancer cell mechanics by mediating cytoskeletal organization and focal adhesion maturation. Oncotarget, 6(18), 15966–15983. 10.18632/oncotarget.3862

22. Maninová, M., & Vomastek, T. (2016). Dorsal stress fibers, transverse actin arcs, and perinuclear actin fibers form an interconnected network that induces nuclear movement in polarizing fibroblasts. The FEBS Journal, 283(20), 3676–3693. 10.1111/febs.13836

23. Marks, P. C., Hewitt, B. R., Baird, M. A., Wiche, G., & Petrie, R. J. (2022). Plectin linkages are mechanosensitive and required for the nuclear piston mechanism of three-dimensional cell migration. Molecular Biology of the Cell, 33(12), ar104. 10.1091/mbc.E21-08-0414

24. Marsigliante, S., Vetrugno, C., & Muscella, A. (2016). Paracrine CCL20 loop induces epithelial-mesenchymal transition in breast epithelial cells. Molecular Carcinogenesis, 55(7), 1175–1186. 10.1002/mc.22360

25. Masiello, N. C., Zucker, M. B., Casella, J. F., Flanagan, M. D., & Lin, S. (1982). Effect of cytochalasin D on thrombin-induced actin filaments in platelets. Nature, 298(5870), 202. 10.1038/298202b0

26. Mason, T. G. (2000). Estimating the viscoelastic moduli of complex fluids using the generalized Stokes-Einstein equation. Rheologica Acta, 39(4), 371–378. 10.1007/s003970000094

27. Mendez, M. G., Restle, D., & Janmey, P. A. (2014). Vimentin enhances cell elastic behavior and protects against compressive stress. Biophysical Journal, 107(2), 314–323. 10.1016/j.bpj.2014.04.050

28. Moeendarbary, E., Valon, L., Fritzsche, M., Harris, A. R., Moulding, D. A., Thrasher, A. J., Stride, E., Mahadevan, L., & Charras, G. T. (2013). The cytoplasm of living cells behaves as a poroelastic material. Nature Materials, 12(3), 253–261. 10.1038/nmat3517

29. Montague, T. G., Cruz, J. M., Gagnon, J. A., Church, G. M., & Valen, E. (2014). CHOPCHOP: A CRISPR/Cas9 and TALEN web tool for genome editing. Nucleic Acids Research, 42(Web Server issue), W401-407. 10.1093/nar/gku410

30. Morgan, M. J., & Liu, Z. (2011). Crosstalk of reactive oxygen species and NF-κB signaling. Cell Research, 21(1), 103–115. 10.1038/cr.2010.178

31. Munder, M. C., Midtvedt, D., Franzmann, T., Nüske, E., Otto, O., Herbig, M., Ulbricht, E., Müller, P., Taubenberger, A., Maharana, S., Malinovska, L., Richter, D., Guck, J., Zaburdaev, V., & Alberti, S. (2016). A pH-driven transition of the cytoplasm from a fluid- to a solid-like state promotes entry into dormancy. eLife, 5, e09347. 10.7554/eLife.09347

32. Najafi, J., Dmitrieff, S., & Minc, N. (2023). Size- and position-dependent cytoplasm viscoelasticity through hydrodynamic interactions with the cell surface. Proceedings of the National Academy of Sciences of the United States of America, 120(9), e2216839120. 10.1073/pnas.2216839120

33. Patteson, A. E., Pogoda, K., Byfield, F. J., Mandal, K., Ostrowska-Podhorodecka, Z., Charrier, E. E., Galie, P. A., Deptuła, P., Bucki, R., McCulloch, C. A., & Janmey, P. A. (2019). Loss of Vimentin Enhances Cell Motility through Small Confining Spaces. *Small (Weinheim an Der Bergstrasse,* Germany), 15(50), e1903180. 10.1002/smll.201903180

34. Pegoraro, A. F., Janmey, P., & Weitz, D. A. (2017). Mechanical Properties of the Cytoskeleton and Cells. Cold Spring Harbor Perspectives in Biology, 9(11), a022038. 10.1101/cshperspect.a022038

35. Pollard, T. D. (1976). The role of actin in the temperature-dependent gelation and contraction of extracts of Acanthamoeba. The Journal of Cell Biology, 68(3), 579–601. 10.1083/jcb.68.3.579

36. Prahlad, V., Yoon, M., Moir, R. D., Vale, R. D., & Goldman, R. D. (1998). Rapid movements of vimentin on microtubule tracks: Kinesin-dependent assembly of intermediate filament networks. The Journal of Cell Biology, 143(1), 159–170. 10.1083/jcb.143.1.159

37. Ran, F. A., Hsu, P. D., Wright, J., Agarwala, V., Scott, D. A., & Zhang, F. (2013). Genome engineering using the CRISPR-Cas9 system. Nature Protocols, 8(11), 2281–2308. 10.1038/nprot.2013.143

38. Rotsch, C., & Radmacher, M. (2000). Drug-Induced Changes of Cytoskeletal Structure and Mechanics in Fibroblasts: An Atomic Force Microscopy Study. Biophysical Journal, 78(1), 520–535. 10.1016/S0006-3495(00)76614-8

39. Sanchez, A. D., & Feldman, J. L. (2017). Microtubule-organizing centers: From the centrosome to non-centrosomal sites. Current Opinion in Cell Biology, 44, 93–101. 10.1016/j.ceb.2016.09.003

40. Sarria, A. J., Nordeen, S. K., & Evans, R. M. (1990). Regulated expression of vimentin cDNA in cells in the presence and absence of a preexisting vimentin filament network. The Journal of Cell Biology, 111(2), 553–565. 10.1083/jcb.111.2.553

41. Sbalzarini, I. F., & Koumoutsakos, P. (2005). Feature point tracking and trajectory analysis for video imaging in cell biology. Journal of Structural Biology, 151(2), 182–195. 10.1016/j.jsb.2005.06.002

42. Schmid, J. A., Birbach, A., Hofer-Warbinek, R., Pengg, M., Burner, U., Furtmüller, P. G., Binder, B. R., & de Martin, R. (2000). Dynamics of NF kappa B and Ikappa Balpha studied with green fluorescent protein (GFP) fusion proteins. Investigation of GFP-p65 binding to DNa by fluorescence resonance energy transfer. The Journal of Biological Chemistry, *275*(22), 17035–17042. 10.1074/jbc.M000291200

43. Schuh, M., & Ellenberg, J. (2007). Self-Organization of MTOCs Replaces Centrosome Function during Acentrosomal Spindle Assembly in Live Mouse Oocytes. Cell, 130(3), 484–498. 10.1016/j.cell.2007.06.025

44. Schwarzl, F. R. (1971). Numerical calculation of storage and loss modulus from stress relaxation data for linear viscoelastic materials. Rheologica Acta, 10(2), 165–173. 10.1007/BF02040437

45. Serres, M. P., Samwer, M., Truong Quang, B. A., Lavoie, G., Perera, U., Görlich, D., Charras, G., Petronczki, M., Roux, P. P., & Paluch, E. K. (2020). F-Actin Interactome Reveals Vimentin as a Key Regulator of Actin Organization and Cell Mechanics in Mitosis. Developmental Cell, 52(2), 210–222.e7. 10.1016/j.devcel.2019.12.011

46. Svitkina, T. M. (2020). Actin Cell Cortex: Structure and Molecular Organization. Trends in Cell Biology, 30(7), 556–565. 10.1016/j.tcb.2020.03.005

47. Svitkina, T. M., Verkhovsky, A. B., & Borisy, G. G. (1996). Plectin sidearms mediate interaction of intermediate filaments with microtubules and other components of the cytoskeleton. The Journal of Cell Biology, 135(4), 991–1007. 10.1083/jcb.135.4.991

48. Tojkander, S., Gateva, G., & Lappalainen, P. (2012). Actin stress fibers – assembly, dynamics and biological roles. Journal of Cell Science, jcs.098087. 10.1242/jcs.098087

49. Trevors, J. T. (2011). The Composition and Organization of Cytoplasm in Prebiotic Cells. International Journal of Molecular Sciences, 12(3), 1650–1659. 10.3390/ijms12031650

50. Tseng, Y., Kole, T. P., & Wirtz, D. (2002). Micromechanical Mapping of Live Cells by Multiple-Particle-Tracking Microrheology. Biophysical Journal, 83(6), 3162–3176. 10.1016/S0006-3495(02)75319-8

51. Wiche, G. (1998). Role of plectin in cytoskeleton organization and dynamics. Journal of Cell Science, 111(17), 2477–2486. 10.1242/jcs.111.17.2477

52. Wirtz, D. (2009). Particle-tracking microrheology of living cells: Principles and applications. Annual Review of Biophysics, 38, 301–326. 10.1146/annurev.biophys.050708.133724

53. Xie, J., Najafi, J., Le Borgne, R., Verbavatz, J.-M., Durieu, C., Sallé, J., & Minc, N. (2022). Contribution of cytoplasm viscoelastic properties to mitotic spindle positioning. Proceedings of the National Academy of Sciences of the United States of America, 119(8), e2115593119. 10.1073/pnas.2115593119

54. Xu, K., Schwarz, P. M., & Ludueña, R. F. (2002). Interaction of nocodazole with tubulin isotypes. Drug Development Research, 55(2), 91–96. 10.1002/ddr.10023

55. Zhang, Y., Zhao, S., Li, M., Li, Y., Feng, F., Cui, J., Xue, Y., Jin, X., & Jiu, Y. (2021). Host cytoskeletal vimentin serves as a structural organizer and an RNA-binding protein regulator to facilitate Zika viral replication. 10.1101/2021.04.25.441301

56. Zhou, H., Xu, J., Zhang, C., & Wen, Y. (2019). Aberrant histone deacetylase 1 expression upregulates vimentin expression via an NF-κB-dependent pathway in hepatocellular carcinoma. Oncology Letters. 10.3892/ol.2019.10309

57. Zimmerman, S. B., & Minton, A. P. (1993). Macromolecular crowding: Biochemical, biophysical, and physiological consequences. Annual Review of Biophysics and Biomolecular Structure, 22, 27–65. 10.1146/annurev.bb.22.060193.000331

